# Spatial transcriptomics reveals unique molecular fingerprints of human nociceptors

**DOI:** 10.1101/2021.02.06.430065

**Authors:** Diana Tavares-Ferreira, Stephanie Shiers, Pradipta R. Ray, Andi Wangzhou, Vivekanand Jeevakumar, Ishwarya Sankaranarayanan, Anna Cervantes, Jeffrey C Reese, Alexander Chamessian, Bryan A. Copits, Patrick M. Dougherty, Robert W. Gereau, Michael D. Burton, Gregory Dussor, Theodore J. Price

## Abstract

Nociceptors are specialized sensory neurons that detect damaging or potentially damaging stimuli and are found in the dorsal root (DRG) and trigeminal ganglia. These neurons are critical for the generation of neuronal signals that ultimately create the perception of pain. These neurons are also primary targets for acute and chronic pain therapeutics. Single-cell transcriptomics on mouse nociceptors has transformed our understanding of pain mechanisms. We sought to generate equivalent information for human nociceptors with the goal of identifying transcriptomic signatures of nociceptors, identifying species differences and elucidating new drug targets. We used spatial transcriptomics to molecularly characterize transcriptomes of single dorsal root ganglion (DRG) neurons from 8 organ donors. We identified 12 clusters of human sensory neurons, 5 of which are C nociceptors; as well as 1 Aβ nociceptor, 2 Aδ, 2 Aβ and 1 proprioceptor subtypes. By focusing on expression profiles for ion channels, G-protein coupled receptors (GPCRs) and other pharmacological targets, we provide a rich map of drug targets in the human DRG with direct comparison to mouse sensory neuron transcriptomes. We also compare human DRG neuronal subtypes to non-human primates showing conserved patterns of gene expression among many cell types, but divergence among specific nociceptor subsets. Finally, we identify sex differences in human DRG subpopulation transcriptomes, including a marked increase in *CALCA* expression in female pruritogen receptor enriched nociceptors. Our data open the door to development of drug discovery programs for new pain targets and unparalleled molecular characterization of clinical sensory disorders.

**One Sentence Summary:** We used spatial transcriptomics to molecularly characterize human sensory neurons, comparing them to mouse and non-human primate finding similarities but also divergence, in particular among drug targets.

## INTRODUCTION

Pain is a major medical problem that has been treated for millennia with drugs whose origins can be traced to natural products (1). While some new mechanism-based therapeutics have recently been approved for treatment of pain, these were developed based on biochemical observations in clinical studies, such as the calcitonin gene-related peptide (CGRP) link to migraine headache (2). There has been an unsatisfying failure to translate preclinical work on peripheral pain mechanisms, which has largely been done in rodents, into effective pain therapeutics (3, 4). A potential explanation for this failure to translate is that important species differences in nociceptor molecular phenotypes exist between mice and humans, an idea partially supported by bulk RNA sequencing experiments (5, 6), and other lines of evidence (7, 8). Nociceptors are the first neurons in the pain pathway and express a broad variety of receptors that allow them to respond to stimuli arising from the environment, from local cells native to tissues, and from infiltrating immune cells that may be involved in inflammation or other processes (9–12). These neurons increase their excitability in both acute and chronic pain states, and changes in their excitability phenotype, such as the generation of spontaneous activity, are directly linked to chronic pain states like neuropathic pain (13). Therefore, nociceptors are excellent target cells for acute and chronic pain drugs that are badly needed to improve pain treatment. In the work described here, we have created a high-resolution map of human sensory neurons in the DRG, including nociceptors, with the goal of accelerating discovery and/or validation of high-quality drug targets that can be manipulated to improve pain treatment.

Single cell sequencing of DRG neurons has delineated the molecular architecture of somatosensory neuron subtypes in the mouse (14–16), elucidated their developmental transcriptional paths (17) and characterized how these neurons change phenotype in response to injury (18, 19). However, it is not clear how this information can be applied to humans because a corresponding transcriptomic map of human sensory neurons does not exist. Most contemporary single cell profiling studies use nuclear RNA sequencing because this technology is scalable, fully commercialized and widely available (20). However, human DRG neurons are among the largest in the body (20 – 100 µm diameter) (21) and also have large nuclei, creating challenges for many sequencing platforms. Sensory neurons are also postmitotic cells with large cytoplasmic volumes that contain a high concentration of extra-nuclear RNA. Sequencing technologies that combine spatial resolution with the ability to accurately sample cytoplasmic RNA may reveal a clearer picture of the full neuronal transcriptome (22) which is important when looking for drug targets that may have low expression. To overcome these issues and fill this gap in knowledge with respect to human sensory neuron transcriptomes, we have conducted spatial sequencing experiments on human, lumbar dorsal root ganglia (DRG) obtained from organ donors. We identified one proprioceptor, two Aß low-threshold mechanoreceptors (LTMRs), one Aß nociceptor, one Aδ-LTMR, one Aδ high-threshold mechanoreceptor (HTMR), one C-LTMR and five nociceptor subtypes. We have compared our findings to both mouse (16) and non-human primate datasets (23), finding many similarities, but also important differences, many of which have important implications for pain target identification. Because sex differences in pain mechanisms are increasingly recognized (24, 25), we performed our studies with an equal number of male and female samples. We anticipate that our data will advance our understanding of molecular pain mechanisms in humans and create a new path forward for pain and itch therapeutic development (4).

## RESULTS

### Spatial transcriptomics generates near single-neuron resolution

We generated whole cell transcriptomes for single neurons, using the 10x Genomics Visium Spatial Gene Expression platform (26, 27). This technology uses 55 µm barcoded-spots printed on the capture area of Visium slides. Human DRGs, collected within 4 hrs of cross-clamp from neurologically dead organ donors (4 female and 4 male, details on organ donors provided in **Table S1**), were sectioned into the capture areas of the Visium slides, stained and imaged (Figure 1A). After tissue permeabilization, mRNA from each section was bound to barcoded primers and subsequently processed for library preparation and RNA sequencing. We obtained on average ∼52M reads and detected an average total of ∼24,000 genes per section, for a total of ∼830M reads from 16 tissue sections (**Figure S1A**). Because each section was stained and imaged, the barcoded mRNAs and respective genes’ location can be visualized within each DRG section using Loupe Browser (10x Genomics). Additionally, barcoded spots can be selected based on their position in the tissue (**Figure S1B**). To generate near single-neuron resolution, we selected all barcodes that overlapped a single neuron in all sections and processed them for downstream analysis. From two tissue sections from each donor (total 16 sections), we identified 4,356 barcodes that overlap a single neuron (‘neuronal barcodes’) and 12,118 barcodes that directly surround neurons (’surrounding barcodes’). The remaining 20,725 barcoded spots were classified as ‘other barcodes’. Barcodes that overlapped multiple neurons were excluded. We optimized tissue permeabilization to enhance neuronal RNA elution onto the slides to develop neuronally-enriched libraries (**Figure S2**). We detected a higher number of RNA molecules, and a higher number of unique genes in the neuronal barcodes (**Figure S1C**). In addition, neuronal barcodes had a distinct gene expression profile from surrounding and other barcodes (**Figure S1D**).

**Figure 1:**
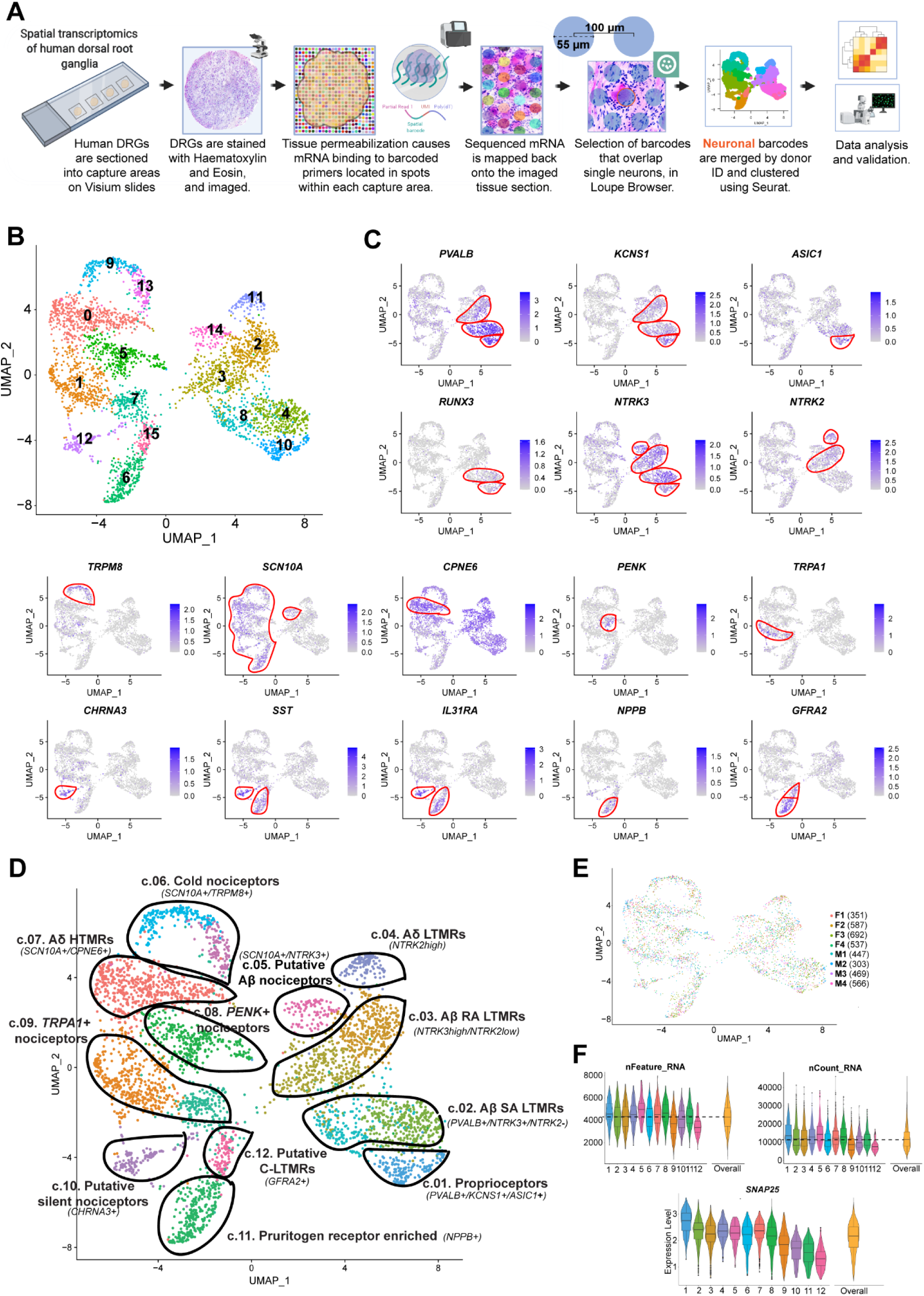
Identification of neuronal subtypes in human DRG using spatial transcriptomics. **(A)** Overview of the workflow and analysis. **(B)** UMAP plot showing the 16 clusters generated by Seurat’s workflow. **(C)** UMAP plots of the expression of gene markers that were used to label neuronal clusters. **(D)** UMAP plot showing the 12 labeled human DRG neuronal clusters that were curated from the original 16 clusters, which are still shown with color coding matching panel **(B)**. Neuronal barcodes (barcoded spots that overlap single neurons) were manually selected in Loupe Browser and clustered using Seurat package in R. **(E)** UMAP plot shows the contribution of each donor for cluster formation. The number of barcodes per donor used for clustering are in parenthesis. **(F)** Violin plots show consistent distributions of the number of detected genes (nFeature_RNA), the counts of unique RNA molecules (nCount_RNA), and the average expression for the neuronal marker *SNAP25* across clusters. The numbers on the x-axis correspond to cluster numbers.

Neuronal barcodes with both a low number of reads and a low count for the neuronal marker *SNAP25* were removed, as described in methods. A total of 3,952 neuronal barcodes were grouped by donor ID and clustered using Seurat’s anchor integration workflow followed by graph-based clustering (28) (see Methods for detailed information and **Figure S3**). Initially, Seurat generated 16 clusters (Figure 1B). We highlight several known neuronal markers from the literature that were enriched in these clusters to characterize these subsets of human DRG neurons based on their specific gene enrichment (Figure 1C). We ultimately selected 8 clusters for merging. Each of these were neighboring clusters with highly overlapping gene expression where 2 clusters were merged into 1. This led to 12 final clusters of human DRG neurons (Figure 1D), which are described in detail below. For data quality purposes, we verified that each individual donor contributed neurons to each cluster and that no individual donor was responsible for any particular cluster (Figure 1E). The number of genes and unique RNA molecules detected per cluster as well as the average expression distribution of the neuronal marker *SNAP25* across clusters is shown in Figure 1F.

### Defining the transcriptomes of human sensory neuron subtypes

DRG neurons are derived from neural crest cells and are responsible for transmitting all somatosensation (touch, proprioception, nociception and temperature) from the body to the brain (29). These neurons have been grouped into two main classes based on the diameter of the cell body and the conduction velocity – A and C fibers. Myelinated Aβ-fiber neurons are mostly large diameter cells that innervate the skin through terminal organs that are responsible for detection of non-noxious stimuli, in particular light touch (30, 31). Proprioceptors innervate muscle and other structures and are responsible for communicating signals about the location of our limbs in space. Unmyelinated, small diameter C-fiber neurons are critical for the detection of most noxious stimuli. A∂ neurons are lightly myelinated and have larger diameter than C-fibers but also respond to stimuli in the noxious range. These classes of sensory neurons differentially express specific neurotrophic receptors during development, and into adulthood (29).

Within the A-fiber group, we identified 6 subtypes in the human DRG. The first cluster was classified as proprioceptors based on the expression of *PVALB, NTRK3* and *ASIC1* and depletion of *NTRK2* (*32*). This cluster was also enriched for *KCNS1* and a displayed enriched expression of *RUNX3,* which plays an evolutionarily conserved role in vertebrates in suppressing *NTRK2* in A-fiber proprioceptors (33) (Figure 2A). Aβ slowly adapting (SA) LTMRs innervate hairy and glabrous skin and terminate on Merkel cells (34, 35). These neurons were enriched in *NTRK3* and depleted from *NTRK2,* a pattern of expression consistent with Aβ slowly-adapting (SA) LTMRs in the mouse (36). These neurons were also enriched in *RAMP1* expression, a receptor component for the calcitonin gene-related peptide receptor (CGRP). The end-organs of Aβ rapidly adapting (RA) LTMRs are Meissner and Pacinian corpuscles in glabrous skin and lanceolate endings in hairy skin (35). The Aβ RA LTMR subgroup was likewise identified by expression of *NTRK3* and a low level of expression for *NTRK2* (*36, 37*). Aδ-LTMR are also known as D-hair afferents and terminate as longitudinal lanceolate endings in hair follicles (35). Aδ-LTMR were characterized by their high level of expression of *NTRK2* and lack of *NTRK3* (36, 37). Mice lacking this subset of *Ntrk2* positive neurons are less sensitive to touch and non-responsive to mechanical stimulation after injury (38). This suggests that Aδ fibers may be involved in the development of mechanical allodynia. Aδ fibers have previously been characterized in human skin nerves as similar to “down-hair” Aδ neurons in other species (39). One group of A-fiber neurons expressed both *NTRK3* and *SCN10A*, a voltage-gated sodium channel that is enriched in nociceptive neurons (40). Therefore, we identified this cluster as putative Aβ-nociceptors. Aβ-fibers that respond to noxious stimuli have been reported in other species (41) including monkeys (42). A recent study has demonstrated that humans also have Aβ-fiber nociceptors with nociceptive properties (43). The final cluster of A-fibers had high expression of *NTRK1*, *CPNE6* and *SCN10A,* which is consistent with Aδ high-threshold mechanoreceptors (HTMRs) in the mouse and macaque (14, 44). This cluster also expressed *CALCA* and *LPAR3*.

**Figure 2:**
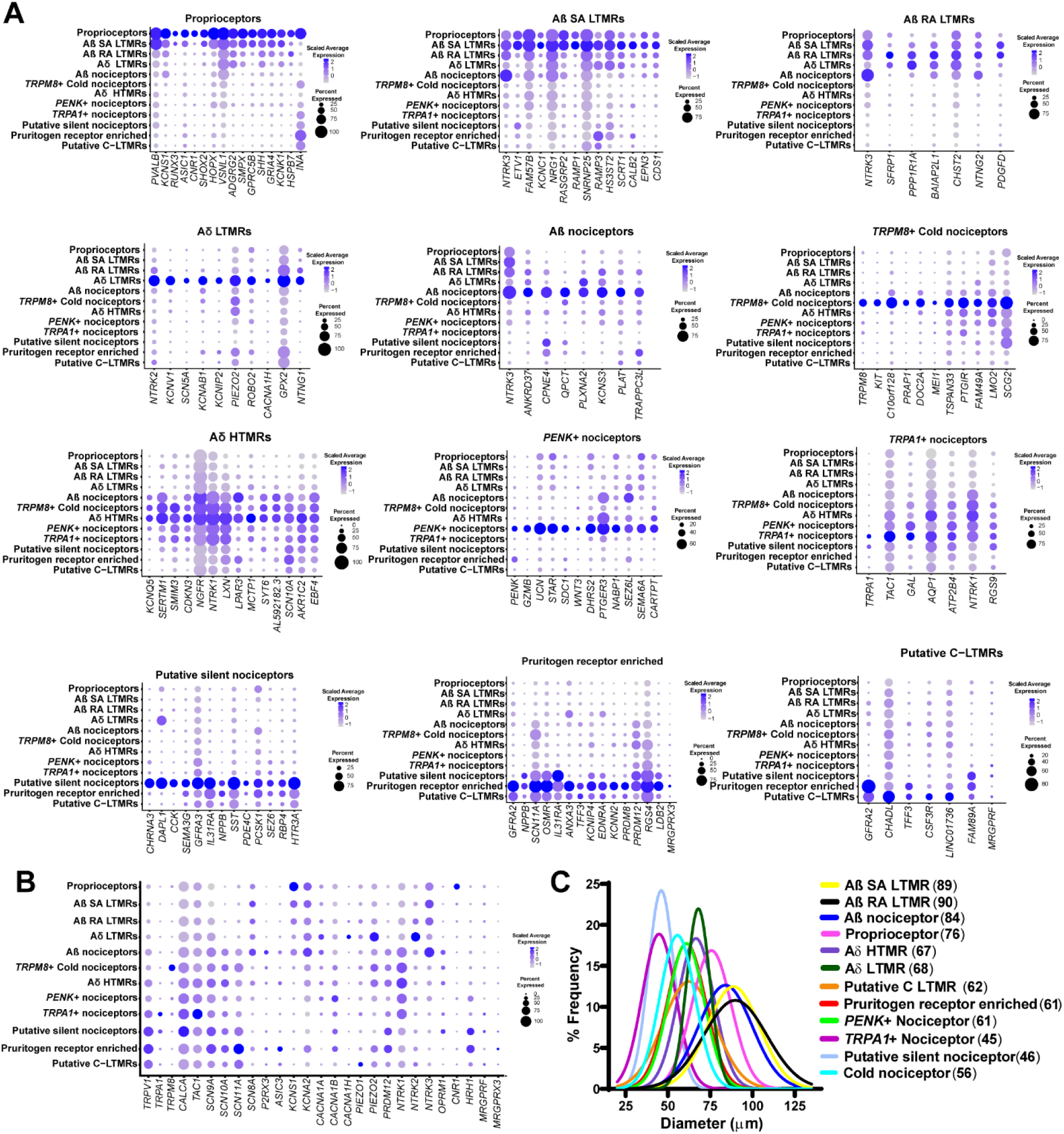
Enriched gene expression in human DRG neuronal clusters and spatial visualization of neuronal subtypes. **(A)** Dot-plots showing the top genes for each neuronal sub-population and how they are expressed across all clusters. The size of the dot represents the percentage of barcodes within a cluster and the color corresponds to the average expression (scaled data) across all barcodes within a cluster for each gene shown. **(B)** Dot-plot showing the expression of known pain genes and markers across clusters. **(C)** Neuronal clusters were mapped back to DRG sections to visualize neurons within the DRG. Diameters of neurons with visible nuclei were measured to ascertain and plot cell sizes for each cluster with mean diameter (in µm) shown in parenthesis. Gaussian curve fits are shown for visualization purposes.

We also identified 5 subtypes of C-fiber nociceptors and a putative C-LTMR cluster (Figure 2A). *TRPM8*, a known menthol and cold sensitive channel, labelled the cold nociceptors (45). This cluster expressed *SCN10A* but little *TRPV1,* a unique feature compared to other human nociceptor clusters. *PENK,* an endogenous opioid and precursor to several enkephalins (46), was enriched in another C nociceptor cluster. This cluster also uniquely expressed the peptide transmitter gene *UCN*, encoding urocortin, and was enriched for the prostaglandin E_2_ (PGE_2_) receptor PTGER3, encoding the EP3 receptor that is distinct among PGE_2_ receptors in producing analgesia upon agonist binding (47). Another cluster of C-fibers was distinguished by *TRPA1* expression. This sub-population also showed very high expression for *TAC1* (substance P) and *CALCA* (*48*) even though these neuropeptides were broadly expressed by all nociceptor clusters. This difference in neuropeptide expression is an important distinction between human and rodent sensory neurons, likely indicating that peptidergic and non-peptidergic subsets of sensory neurons do not exist in humans (7, 8, 49).

The specific expression of *CHRNA3* identified a cluster of putative ‘silent’ nociceptors (50) (Figure 2A). ‘Silent’ nociceptors correspond to a subset of C-fibers that innervate joints, viscera and skin and are often referred to as mechano-insensitive C-fibers (CMi). They are unresponsive to noxious mechanical stimuli under normal conditions, but are sensitized and become mechanically sensitive after inflammatory stimulation, and likely play key roles in certain pain disorders (50–53). The silent nociceptor cluster expressed a large array of ion channels including the serotonin receptor *HTR3A*; purinergic receptors *P2RX4, P2RX3, P2RX6* and *P2RX7*; proton receptor *ASIC3*; and glutamate receptors *GRIK2, GRIK3, GRIK4, GRIK5, GRID1, GRIN1, GRIA3,* and *GRIA4* (**File S1**), which may underlie the sensitivity of this subset of neurons to inflammatory mediators. These neurons also expressed the H1 histamine receptor gene, *HRH1*, which is known to sensitize these neurons to mechanical stimulation, and is also a likely pathway for histamine-induced itch in humans (54, 55). Therefore, this subset of C-fibers likely also participates in the generation of itch signals from the periphery. A separate pruritogen receptor enriched cluster was classified based on the expression of *NPPB*, *GFRA2* and *IL31RA* (*56*), although these latter 2 genes were also found in other populations. Our data also shows that *SCN11A* has a very high expression level in this sub-population. Nav1.9 (*SCN11A*) gain of function mutations can lead to congenital insensitivity to pain (CIP) or partial loss of pain sensation. Studies in mice have reported that the mutation causes a pruritic phenotype (57, 58). Humans with Nav1.9 mutations report a severe pruritis (57, 59). Mechanisms associated with the enrichment of *SCN11A* in itch nociceptors may explain this phenotype. A final C-fiber cluster was enriched in *GFRA2,* a characteristic marker of C-LTMRs in mice (14), and was classified as putative C-LTMRs. This cluster had high similarity in terms of gene expression with the pruritogen receptor enriched population but had lower expression of *NPPB*, a marker for itch nociceptors in mice (**Figure S4A,B**). A distribution of genes associated with pain across human DRG neuronal subtype clusters is shown in Figure 2B. Ranked gene expression by gene for all 12 A-and C-fiber clusters are given in **File S1**.

### Spatial visualization of neuronal subtypes

Lumbar DRG neuronal subtypes did not show any clear spatial organization in any analysed tissue sections. However, we did use visualization of barcode position in DRG sections to measure neuron diameter associated with each of the 12 clusters (see Methods). This independent measure validates that Aβ clusters correspond to the largest diameter neurons in the DRG, while C nociceptors clusters were the smallest (Figure 2C**; Figure S5**). A∂ clusters were intermediate in size between Aβ-and C-fiber neurons, in line with cell size distributions in all other species where this has been assessed (8, 60–62).

### Validation of spatial transcriptome-defined subtypes with RNAscope

Our spatial transcriptomic approach provides detailed insight into the types of neurons present in the human DRG, but there are limitations, such as the lack of pure single neuronal transcriptomes for any given barcode. We have previously demonstrated that RNAscope *in situ* hybridization technology offers highly sensitive detection of neuronal mRNAs in human DRG (8). As a validation tool, we conducted RNAscope experiments on human DRG tissue sections for several mRNAs that showed high abundance in specific neuronal clusters: *PRDM12, NPPB*, *SST, NTRK1-3, PVALB, LPAR3, PENK, TRPM8*. We assessed their co-expression with nociceptor-enriched genes *SCN10A*, *TRPV1*, and *CALCA* (Figure 3A). The nociceptor population (*SCN10A+*, *TRPV1+*, or *CALCA+)* comprised ∼60-70% of all human sensory neurons and were small in diameter (average = 54 µm) (Figure 3B-C). *PRDM12,* a gene that is essential for human pain perception (63) was expressed in ∼74% of DRG neurons and co-expressed *CALCA* (Figure 3D). *CALCA* mRNA was detected in all neuronal clusters and surrounding/other barcodes in the Visium data, likely because *CALCA* mRNA localizes to axons (64) explaining its wide-spread detection. Smaller subdivisions of nociceptors such as the putative silent and pruritogen receptor enriched nociceptor populations (*NPPB*+ or *SST*+) amounted to ∼30% of the population and co-expressed *SCN10A* (Figure 3E-F). *NTRK1,* which is most abundant in the nociceptor clusters, was found in 68% of the neuronal population and co-localized with *SCN10A* (Figure 3G). *NTRK2* which was enriched in the Aδ LTMR cluster, a cluster that is depleted of *SCN10A,* was detected in medium sized neurons (Figure 3C) and showed little co-expression with *SCN10A* (Figure 3H). The proprioceptor, Aβ-LTMR and Aβ-nociceptor marker, *NTRK3*, was found in larger sized neurons and showed slightly higher co-expression with *SCN10A* than *NTRK2*; most likely due its presence in the *SCN10A*+ Aβ-nociceptor cluster (Figure 3I). In the VISIUM dataset, *LPAR3* was enriched in the Aδ HTMR and Aβ nociceptor clusters, but was also lowly expressed in other nociceptor clusters, all of which express *TRPV1*. Similarly, *LPAR3* was expressed in 80% of all sensory neurons, the majority of which were TRPV1-positive (**Figure S6A-C**). *PVALB*, which was highly enriched in the Proprioceptor and Aβ SA LTMR clusters, was found in ∼45% of sensory neurons, half of which were *TRPV1*-negative (**Figure S6A-C**). The cold-nociceptor cluster marker, *TRPM8*, was found in ∼50% of sensory neurons while *PENK*, which was enriched in a different cluster (*PENK* nociceptors), was found in ∼35% of sensory neurons (Figure **S6B-E**). Similar to VISIUM, these two genes did show some overlap (21.2%) using RNAscope but were also detected in separate populations (**Figure S6B-E**). We have previously reported that *TRPV1* mRNA is more widely expressed in human nociceptors than in mouse (8) and *TRPV1* was detected in all nociceptor clusters with Visium spatial sequencing, with the exception of cold nociceptors where it was expressed at very low levels. Using *SCN10A* as a nociceptor marker, we again observed that *TRPV1* was found in most nociceptors (Figure 3J). We next determined if these neurons were functionally responsive to the TrpV1 ligand, capsaicin. Application of capsaicin depolarized all small-sized, dissociated human DRG neurons and caused action potential firing in 75% (Figure 3K). We conclude that RNAscope, spatial sequencing and functional analysis support broad expression of TrpV1 in human nociceptors. As a final validation, previously published RNAscope findings substantiate the proposed neuronal subclusters from Visium sequencing (Figure 3L)(8). For example, we previously proposed *KCNS1* as a marker of human Aβ neurons due to its expression in large-sized neurons that were negative for *CALCA* and *P2RX3* (8). *KCNS1* was also enriched in Aβ clusters using the spatial transcriptomic approach.

**Figure 3.**
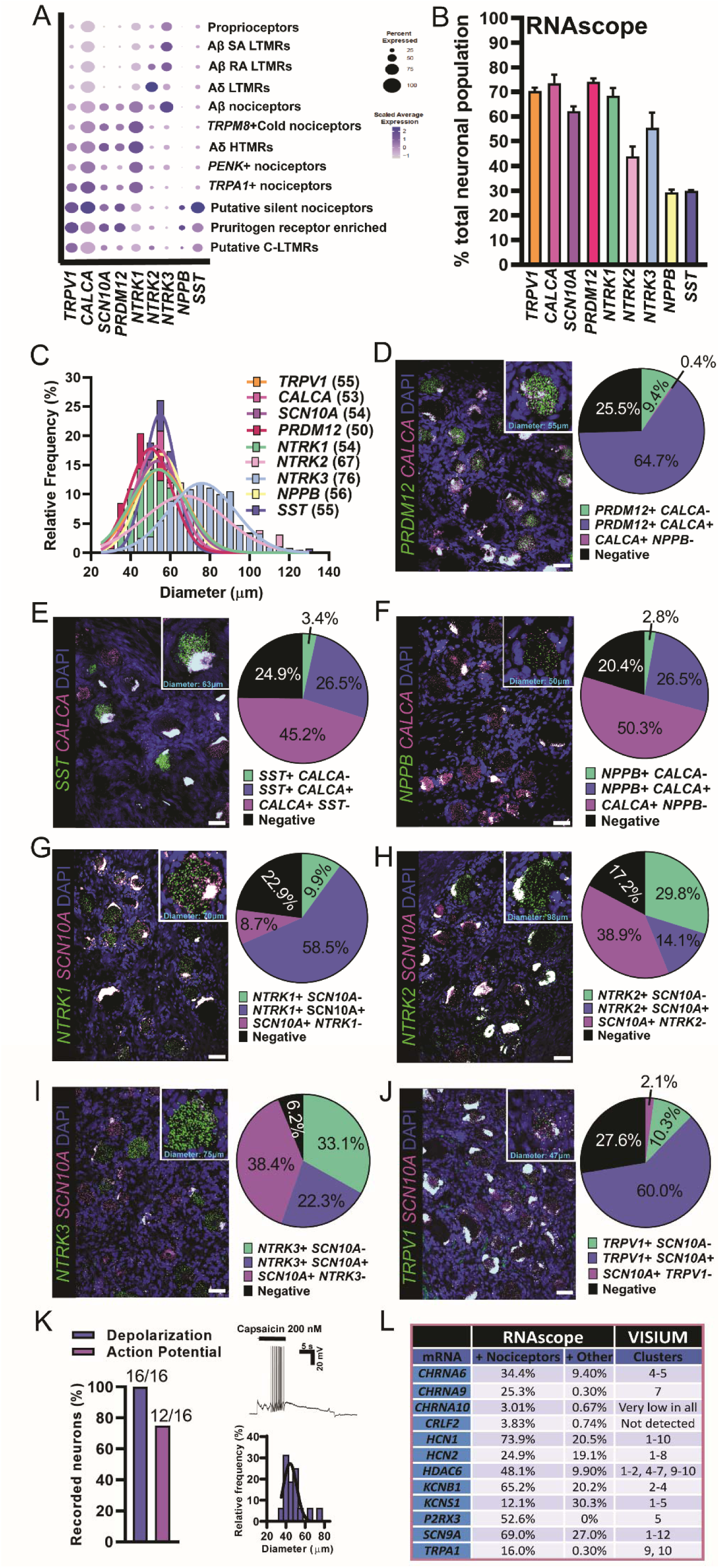
RNAscope *in situ* hybridization and functional validation on human DRG. **(A)** Visualization of Visium gene expression for markers that were used for RNAscope analysis. **(B)** The percentage of neurons expressing each target compared to the total neuronal population. **(C)** The size distribution of all target-positive neurons. Gaussian mean in μm diameter in parentheses. **(D-J)** Merged image for the target of interest (green) with a nociceptor marker (magenta) and DAPI (blue) is shown (scale bar = 50 μm). Inset for each panel shows a blow up of a single neuron. Population distribution of each neuronal marker is shown in the pie chart. **(K)** The TRPV1 agonist, capsaicin (200 nM), was applied to small diameter human DRG neurons *in vitro* causing depolarization (100%) and action potential firing (75%). **(L)** RNAscope data is summarized from (8) and compared to findings from Visium sequencing. The neuronal cluster for each target is listed. Clusters - 1: Proprioceptors, 2: Aβ SA LTMR, 3: Aβ RA LTMR, 4: Aδ LTMR, 5: Aβ nociceptors, 6: *TRPM8*+ Cold nociceptors, 7: Aδ HTMR, 8: *PENK*+ nociceptors, 9: *TRPA1*+ nociceptors, 10: Putative silent nociceptors, 11: Pruritogen receptor enriched, 12: Putative C-LTMRs.

### Sex differences in human sensory neurons

Molecular differences between male and female sensory neurons have been reported in defined population cell sequencing experiments in rodents (65) and inferred from bulk RNA-seq on human DRGs (66), but nothing is known about sex differences in neuronal gene expression in the human DRG. First, it was apparent that males and females have the same DRG neuronal subtypes, because neuronal barcodes from both sexes were clearly represented in all clusters (Figure 4A). We then looked for sex differences within the overall population of neuronal barcodes, and within each specific cluster. With the spatial sequencing approach, neuronal barcodes include mRNA from surrounding cells. To overcome detection of generic sex differences contributed by other cell types, we performed statistical tests on surrounding barcodes (overall surrounding barcodes and specific to each neuronal cluster). We considered genes to be differentially expressed (DE) specifically in neurons if they were not DE in the respective surrounding barcodes (**Figure S7**). Similar to findings in the mouse where sex differences in the neuronal population were small (65), we identified only 44 genes with sex-differential expression in the neuronal barcodes pooled together by sex (Figure 4B and **File S2**). However, this approach pools together expression data for transcriptomically diverse neurons, creating variation that is a product mostly of different cellular phenotypes. To overcome this issue, we looked at potential sex differences in gene expression within each neuronal subtype. Here we found more neuronally-enriched DE genes (Figure 4C, **Files S3-14 for neuronal barcodes, Files S15-27 for surrounding barcodes**). The pruritogen receptor enriched population had the highest number of DE genes (96), suggesting potential molecular differences in mechanisms of pruritis between men and women. We performed gene-set enrichment analysis for DE genes in the itch population using GO Enrichment Analysis resource, PANTHER (67). Several genes were involved in the development of the nervous system and in the response to external stimuli (**File S28**). A striking difference was the increase in *CALCA* expression, which encodes the CGRP protein, found in female pruritogen receptor enriched neurons (Figure 4D). This finding was validated in RNAscope experiments examining *CALCA* expression in *NPPB*-positive neurons from male and female organ donors (Figure 4E**; Figure S8**).

**Figure 4.**
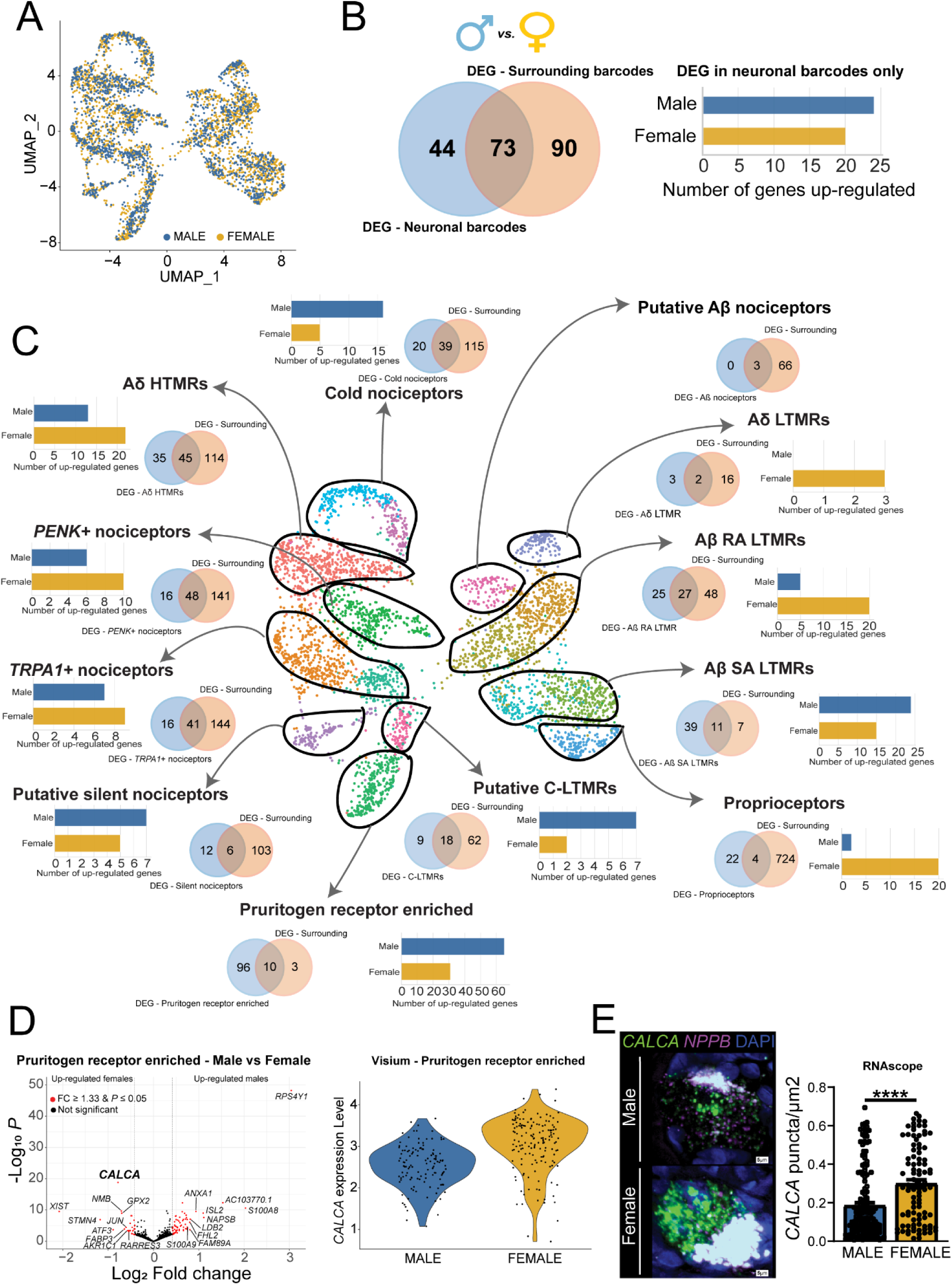
Sex differences in gene expression within human DRG neuronal populations. **(A)** UMAP showing that male and female barcodes are equally represented in all clusters. **(B)** Venn diagram showing the overlap between the number of DE genes in the overall neuronal population (blue, left) and the overall surrounding population of barcodes (beige, right). Bar-plot shows the number of up-regulated genes per sex after removing genes that were also DE in surrounding barcodes. **(C)** Venn diagrams show the overlap between the number of DE genes in each neuronal subtype (blue, left) and the respective surrounding population (beige, right). Bar-plots show the number of up-regulated genes per sex in each cluster after removing genes that were also DE in the respective surrounding barcodes. **(D)** Volcano plot shows DE genes in the pruritogen receptor enriched population after removing DE genes in surrounding barcodes (we highlighted the top 10 genes in each sex ranked by log_2_ fold change). Violin plot shows *CALCA* expression in individual barcodes in males and females within the pruritogen receptor enriched population. **(E)** RNAscope for *CALCA* mRNA colocalized with *NPPB*, a marker of pruritogen receptor enriched nociceptors, with quantification of differences in expression between male and female neurons for amount of *CALCA* expression in the dot-plot. Representative image scale bar = 5 µm. DEG = differentially expressed gene. **** p < 0.0001.

### Similarities and differences between human and mouse DRG neurons with a focus on pharmacological targets

Next, we examined expression of individual genes within gene families, such as ion channels, GPCRs and tyrosine receptor kinases, that are involved in transduction of nociceptive signals by nociceptors and are considered important pharmacological targets for existing or novel drugs. We made comparisons between our spatial transcriptomic dataset from human DRG and mouse single neuron data from DRG that is publicly available at mousebrain.org (16). Most pre-clinical studies are conducted in rodents (in particular, mice) so the comparative expression maps that follow can be used to directly assess similarities and differences in sensory neuron gene expression profiles between mouse and humans.

Voltage-gated sodium channels (VGNaCs) are the foundation of the ability of neurons to carry action potentials and sensory neurons express a unique subset of these genes (68). We observed that VGNaC genes have very similar expression patterns in human and mouse (Figure 5A), demonstrating that the expression of α subunits that encode the pore-forming unit of the channel is conserved. An exception among this family was the *SCN4B* gene which encodes the β4 subunit of the VGNaC. This β subunit is critical for resurgent currents that are key contributors to excitability (69, 70). In mouse *Scn4b* was found mostly in A-fiber neurons, consistent with previous studies (69, 70), yet in human *SCN4B* mRNA was distributed among all sensory neuron types. Since β4 subunits regulate resurgent currents through Nav1.8 channels (71), and these two genes are more highly co-expressed in human nociceptors, this could potentially contribute to enhanced resurgent Nav1.8 currents in those cells, a hypothesis that could be tested in future experiments.

**Figure 5:**
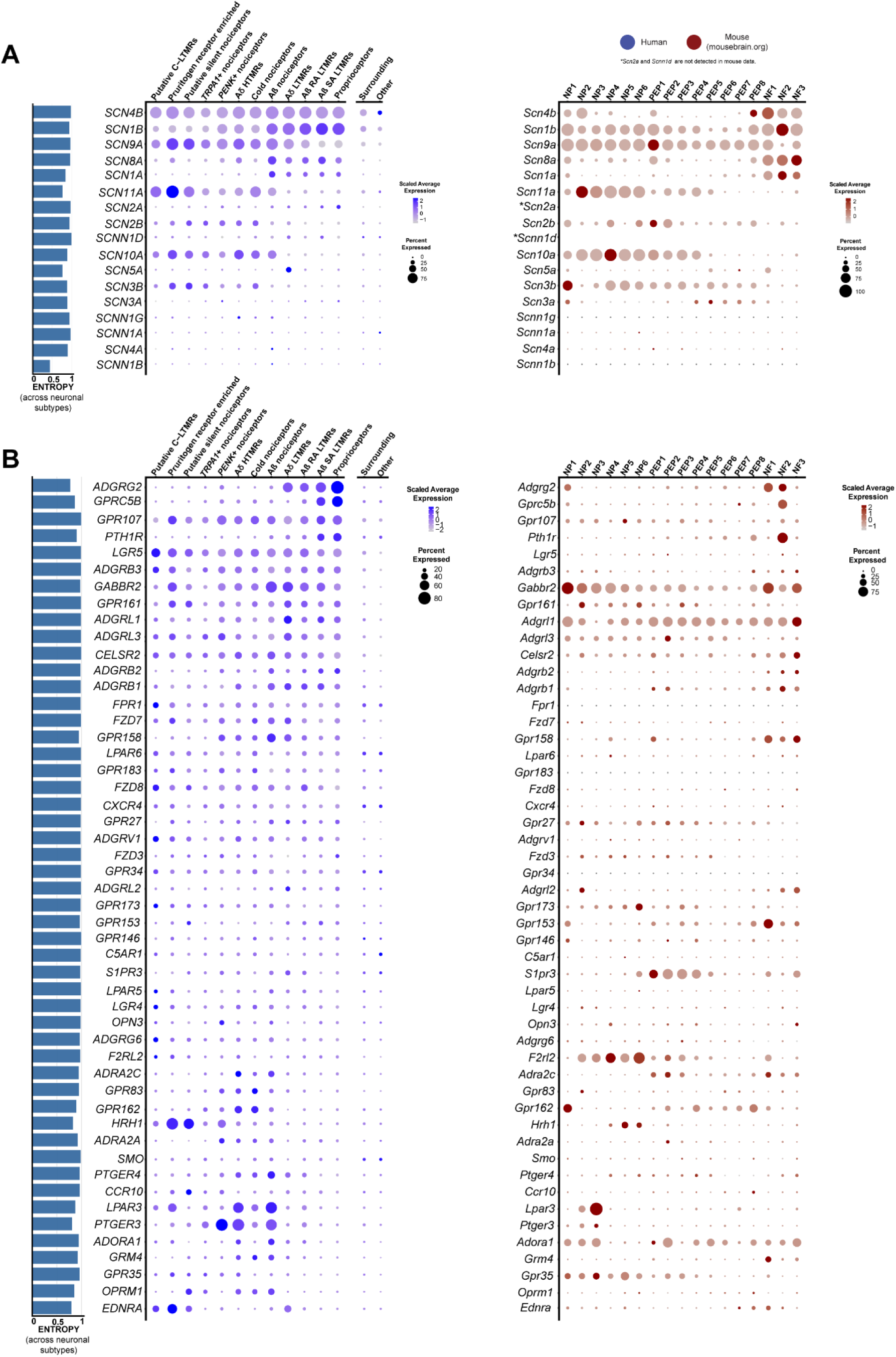
Expression of VGNaC channel and GPCR genes in human and mouse datasets. **(A)** Dot-plots showing the expression of VGNaC channel genes in human spatial transcriptomic (in blue) and mouse single-cell experiments (in red). **(B)** Dot-plots showing the expression of GPCR genes in human spatial transcriptomic (in blue) and mouse single-cell experiments (in red). The size of the dot represents the percentage of barcodes within a cluster and the color corresponds to the average expression (scaled data) across all barcodes within a cluster for each gene shown. Normalized entropy was used as a measure of “specificity of neuronal subtype”, where a score of 0 means a gene is specific to one neuronal subtype and 1 means that a gene has uniform distribution across neuronal subtypes.

G-protein-coupled receptors (GPCRs) are the largest family of receptors in the mammalian genome and have diverse roles in nociceptors ranging from inflammation detection to cell adhesion. These receptors are also important targets for therapeutic development. We compared the expression level and distribution of the top 50 most highly expressed GPCRs in human DRG to their homologs in mouse. While some GPCRs showed consistent patterns of expression, many were divergent suggesting important differences in expression across species for this family of receptors. Two notable differences were the *PTGER3* and *LPAR3* genes (Figure 5B). While *PTGER3* was enriched in the *PENK*+ nociceptor population in humans, it was also expressed by several other nociceptor subtypes while it was restricted to a subset of non-peptidergic neurons in mice. Given the potential for this prostaglandin receptor as an antinociceptive target, this could be important for therapeutic purposes with EP3 agonists (47). *LPAR3* is a receptor for lysophosphatidic acid and has been associated with neuropathic pain (72). This GPCR was broadly expressed in nociceptor subtypes in humans but was again restricted to non-peptidergic nociceptors in mice. Receptors of the metabotropic glutamate receptor family (*GRM*) also showed divergent expression across species (Figure 5B), consistent with the previous observation that group I *GRM* family genes are not detected in human DRG (5). Some GPCRs did show strong conservation of expression, for instance *GABBR2*, encoding a subunit of the GABA_B_ receptor complex, which is likely the most highly expressed Gα_i_-coupled receptor in sensory neurons in both humans and mice.

The characterization of expression of interleukins (IL) and their receptors in neuronal subpopulations can reveal how their ligands may interact with different populations of sensory neurons in different species. *IL31RA*, for instance, was more broadly expressed in human DRG neurons than in mouse where the gene was restricted to itch nociceptors (**Figure S9)**, as shown previously using *in situ* hybridization (73). Anti-inflammatory IL receptors, *IL4R*, *IL10RA* and *IL13RA1* showed broader expression across human sensory neuron subtypes than in mice, where *Il4r* was not detected. Other genes such as *IL6ST* showed conserved expression in humans and mice. We examined expression of other gene families across human and mouse neuronal subtypes including: acid sensing ion channels (ASIC) (**Figure S10**), anoctamins (**Figure S11**), aquaporins (**Figure S12**), calcium channels (**Figure S13**), chloride channels (**Figure S14**), cholinergic receptors (**Figure S15**), ionotropic GABA receptors (**Figure S16**), gap-junction/connexins (**Figure S17**), ionotropic glutamate receptors (**Figure S18**), glycine receptors (**Figure S19**), neuropeptide genes (**Figure S20**), potassium channels (**Figure S21**), ionotropic purinergic receptors (**Figure S22**), transient receptor potential channels (**Figure S23**) and transcription factors involved in neuronal differentiation (**Figure S24**). We also looked at the expression of genes that encode for proteins that are part of the understudied druggable genome (74) (**Figure S25-S27**). We detected 56 understudied GPCRs (out of 117) (**Figure S25**), 49 understudied ion-channels (out of 62) (**Figure S26**) and 133 kinases (out of 150) (**Figure S27**) in human DRG neuronal subtypes. Lastly, we created an expression map of the genes with lowest normalized entropy representing genes with the greatest variance of expression across neuronal subtypes (**Figure S28**).

### Comparison of human and non-human primate sensory neuron subtypes

Next, we took advantage of a recently published single cell dataset from non-human primate DRG to compare neuronal subtypes between human and macaque. Distinct orthology was identified between human and macaque subpopulations, but several orthologs had a many-to-one mapping (e.g. between A LTMR in Rhesus and Aδ LTMR and Aβ RA-LTMR in humans), suggesting that some of the macaque subpopulations could be further subdivided into distinct populations. The subpopulation orthology and corresponding gene expression clusters are shown in Figure 6A-C. Comparison of human and mice neuronal subpopulation transcriptomes show many important changes in neuronal DRG expression (8, 49, 75), which are consistent with macaque and mouse differences (23). Based on analysis of the most lineage restricted human DRG genes in Figure 6, expression enrichment was found to be broadly conserved in humans and macaque, but regulatory divergences in some important sensory genes were also observed.

**Figure 6:**
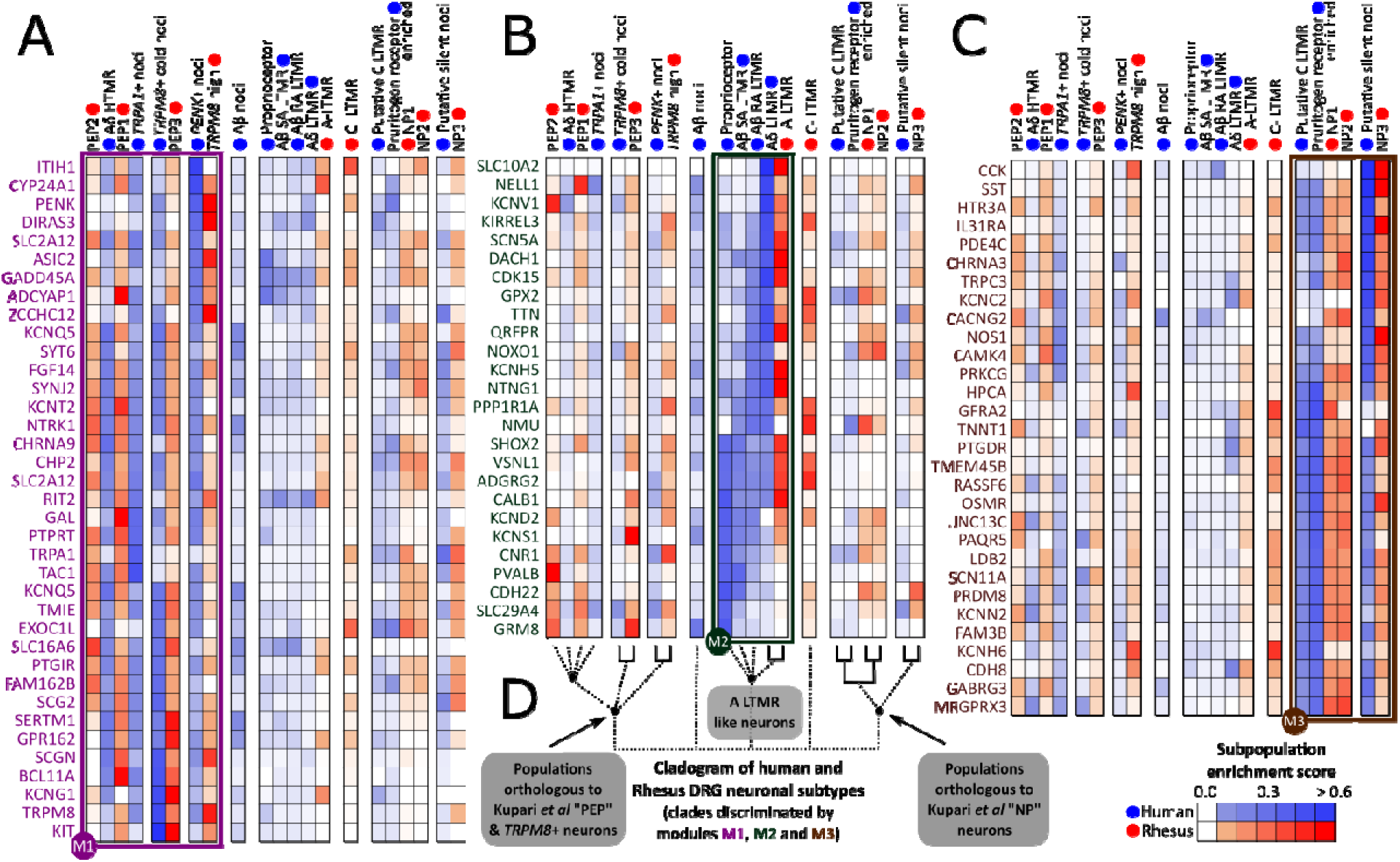
Orthologous neuronal populations between human and macaque. Lineage-restricted human DRG neuronal genes that have high dynamic range of expression in macaque DRG neuronal populations (from Smart-seq2) identify broadly orthologous cell types in the human and macaque, showing evolutionary conservation of gene expression patterns (shown in three gene modules M1, M2 and M3 in **A, B & C** respectively). Expression enrichment analysis show evolutionary divergence in cell types like macaque C-LTMRs and human proprioceptors. Macaque C-LTMRs show increased expression for some human pruritogen receptor enriched neuron and putative C-LTMR genes (*GFRA2*, *KCNH6*, *TMEM45B* in module M3), as well as enrichment for some human A-LTMR genes (*NMU*, *GPX2*, *KIRREL3* in module M2). Macaque PEP2 enriched genes are primarily found in module M1, but a small subset is enriched in human proprioceptors and LTMRs (*PVALB, GRM8, KCNV1* in module M2). **D** shows the orthology among neuronal populations based on the hDRG lineage-restricted genes, with strong orthologies indicated with solid lines.

*PVALB* gene expression in humans was enriched in proprioceptors and Aβ SA-LTMRs, but not in the corresponding macaque LTMR populations (Figure 6B). Instead, it was enriched in macaque peptidergic PEP2 population, which is transcriptionally closer to the human Aδ HTMRs and TRPA1+ nociceptors where *PVALB* is de-enriched. This species difference is striking because *Pvalb* is a marker of A-fiber LTMR neurons in rodents (76). The neuronal calcium sensor *HPCA* (Hippocalcin) was enriched in human pruritogen receptor enriched, silent nociceptors and putative C-LTMRs, and their macaque orthologs (non-peptidergic (NP) subpopulations; Figure 6C). However, in macaques it was additionally enriched in *TRPM8*+ and PEP1 subpopulations but de-enriched in the human orthologs of these populations. Finally, at the population level, macaque C-LTMRs showed enrichment for some human pruritogen receptor enriched neuron and putative C-LTMR genes (*GFRA2, KCNH6, TMEM45B*), as well as enrichment for some human A-LTMR genes (*NMU, GPX2, KIRREL3*) (Figure 6B and C). Human putative C-LTMRs, on the other hand were de-enriched for all of the analyzed human A-LTMR genes. This suggests that macaque C-LTMRs are transcriptionally divergent from all identified human subpopulations, including the human A-LTMRs and putative C-LTMR populations. Markers for Aβ nociceptors are not enriched in specific macaque populations, and hence are not shown (along with some additional markers for human putative C-LTMRs and human pruritogen receptor enriched neurons) in Figure 6. The complete data for all 111 analyzed gene markers can be found in **File S30**.

## DISCUSSION

### Single neuron transcriptomics and the path to improved pain targets

Our work demonstrates that spatial transcriptomics can be used to generate near-single neuron resolution to define, for the first time, molecular profiles of neuronal subtypes in the human DRG. Our findings demonstrate many similarities but also substantial differences between mouse, where most single nociceptor transcriptome work has been done (15, 16, 18). Some of these differences may be explained by technical issues related to sequencing methods, however, our demonstration of more consistent similarities between macaque and human, where different sequencing techniques were also applied (44), makes this possibility less likely. An important outcome of our experiments is the ability to now directly assess target expression across species with single neuron resolution. We lay out these expression profiles for most pharmacologically relevant targets in mouse and human DRG. This expression map can allow investigators to initiate DRG-focused target identification efforts with human neuronal transcriptome insight and then make data-driven choices about model species and testing paradigms that best fit the chosen development pipeline.

### Divergence of human and rodent DRG neurons

An area where evolutionary divergence between mouse and human sensory neurons is most striking is in neuropeptide, *TRPV1* and *NTRK1* expression. In mice and rats these genes are developmentally regulated with expression in all nociceptors in early development and then silencing in specific populations following postnatal target innervation (17, 77, 78). In contrast to rodents, most human nociceptors share expression of these genes suggesting a blending of many of the markers of peptidergic and non-peptidergic nociceptors that are found in other species, in particular the mouse. This indicates that the peptidergic and non-peptidergic nomenclature is unlikely to have utility for describing human nociceptors (7, 8, 49, 79). Our findings suggest that development programs that silence *TPRV1* and *NTRK1* expression in subgroups of nociceptive sensory neurons are not engaged in humans.

We found important differences in receptor and neuropeptide expression in the pruritogen receptor enriched population with more widespread expression of many markers that have been identified in mouse, in particular into the silent nociceptor and putative C-LTMR subtypes. This is consistent with previous studies that have shown species differences in expression of pruritogen receptor genes such as the IL31 receptor (73). It is also consistent with a recent study comparing macaque and human DRG expression of pruritogen receptors *MRGRPD* and *MRGPRX1* that demonstrated co-expression with *TRPV1* (80). This contrasts with mouse experiments where *Mrgprd* is expressed by a subset of neurons that are devoid of *Trpv1* (81). Klein and colleagues also demonstrated that histamine creates a greater area of flare and wheal in human volunteers than MRGPRD or MRGPRX1 agonists (81). This finding is explained by the expansion of neuronal populations that express *HRH1*, the H1 histamine receptor, in human DRG. Interestingly, we do not find expression for most known markers of C-LTMRs in human DRG, including a lack of markers that were recently identified in a nuclear sequencing study from macaques (44), making this population challenging to identify in our study. The exception was *GFRA2* expression which marks C-LTMRs across species (16, 23), enabling putative identification of C-LTMRs in human DRG.

### Moving human DRG transcriptomics to the clinic

Our spatial transcriptomic characterization of human DRG neuronal subtypes should enable new discoveries in the pain and sensory neuroscience field. One advance is the identification of sets of markers that can be used to molecularly phenotype subtypes of sensory neurons that can be sampled through skin biopsies and other methods from neuropathy patients. While there are clear indications of pathology in sensory neurons indicated from clinical skin biopsy studies, these are almost always grouped into small and large fiber neuropathies, but further distinctions are not made. Our work enables greater mechanistic insight from routine clinical tests. The finding that neuronal transcriptomes in the DRG are stable unless frank axonal injury has occurred (18) suggests that our dataset can be utilized for this purpose almost immediately. Our dataset can also be used to mine for pharmacological targets that can be used to specifically manipulate the excitability of different subsets of nociceptors. This offers the possibility for development of pain targets that are identified based entirely on human transcriptomic data. Our dataset is “sex-aware” insofar as it contains both male and female samples. We highlight sex differences (e.g. greater CGRP expression in the pruritogen receptor enriched population in females) that may be important considerations for therapeutic development. Finally, this dataset can be a foundation to more thoroughly vet targets that have been discovered in studies of peripheral nerves in animal pain models. Our findings now make it possible for conservation of gene expression in human nociceptors to be a first step in de-risking pain targets for future drug development (4).

### Study Limitations

The most important limitation is that the spatial transcriptomic approach only approximates single neuron transcriptomes in the human DRG. Combining our findings with nuclear sequencing will improve resolution. We anticipate that these techniques will complement one another to provide a comprehensive picture of human DRG neuronal transcriptomes. While we have sampled from 8 organ donors, our data cannot account for possible difference in gene expression across the human population, or at earlier stages of life. Future studies can use our foundation to address these important questions. Finally, as we have mentioned throughout the manuscript, species comparisons rely on different sequencing techniques, and different post-mortem intervals so this should be considered in interpreting the data reported here.

## MATERIALS AND METHODS

### Study design

All human tissue procurement procedures were approved by the Institutional Review Boards at the University of Texas at Dallas (UTD). Human lumbar dorsal root ganglion tissues were collected from organ donors within 4 hours of cross-clamp and from neurologic determination of death donors. Donor information is provided in **Table S1**. DRGs used for Visium and RNAScope were frozen on dry ice at the time of extraction and stored in a −80°C freezer. The human DRGs were gradually embedded in OCT in a cryomold by adding small volumes of OCT over dry ice to avoid thawing. DRGs used for Visium were cryosectioned onto SuperFrost Plus charged slides at 10µm. To sample a larger subset of neurons, two sections were utilized from each donor and each 10µm section was 200µm apart. DRGs used for RNAscope were sectioned at 20µm. DRGs used for primary neuronal cultures were placed in artificial cerebral spinal fluid (aCSF) over ice at the time of surgical extraction and transported immediately to the University of Texas at Dallas for processing. These tissues were then used for spatial transcriptomics and all downstream data analysis.

### Visium Spatial Gene Expression

Visium tissue optimization and spatial gene expression protocols were followed exactly as described by 10x Genomics (https://www.10xgenomics.com/) using Haematoxylin and Eosin as the counterstain. Optimal permeabilization time was obtained at 12 min incubation with permeabilization enzyme. Imaging was conducted on an Olympus vs120 slide scanner. DRGs from donors 1-8 were used. mRNA library preparation and sequencing (Illumina Nextseq 500) were done at the Genome Center in the University of Texas at Dallas Research Core Facilities.

### Visium Spatial RNA-seq – mapping raw counts and alignment of barcoded spots with imaged sections

The output data of each sequencing run (Illumina BCL files) was processed using the Space Ranger (v1.1) pipelines provided by 10x Genomics. Samples were demultiplexed into FASTQ files using Space Ranger’s mkfastq pipeline. Space Ranger’s count pipeline was used to align FASTQ files with brightfield microscope images previously acquired, detect barcode/UMI counting, and map reads to the human reference transcriptome (Gencode v27 and GRCh38.p10) (82). This pipeline generates, for each sample, feature-barcode matrices that contain raw counts and places barcoded spots in spatial context on the slide image (cloupe files). Gene expression with spatial context can, then, be visualized by loading cloupe files onto Loupe Browser (v4.2.0, 10x Genomics). The space ranger output statistics for raw data can be found in **File S25**.

### RNAscope *in situ* hybridization

RNAscope *in situ* hybridization multiplex version 1 and version 2 were performed as instructed by Advanced Cell Diagnostics (ACD) and as previously described (8). A table of all probes, combinations, and donor tissues used is shown in **Table S2**. All tissues were checked for RNA quality by using a positive control probe cocktail (ACD) which contains probes for high, medium and low-expressing mRNAs that are present in all cells (ubiquitin C > Peptidyl-prolyl cis-trans isomerase B > DNA-directed RNA polymerase II subunit RPB1). A negative control probe against the bacterial DapB gene (ACD) was used to reference non-specific/background label. The donor id # for the DRGs that were used in each experiment is indicated in the Figure captions.

### RNAscope Imaging and Analysis

DRG sections were imaged on an Olympus FV3000 confocal microscope at 20X or 40X magnification. The acquisition parameters were set based on guidelines for the FV3000 provided by Olympus. In particular, the gain was kept at the default setting 1, HV ≤ 600, offset = 4, and laser power ≤ 20%. Large globular structures and/or signal that auto-fluoresced at the 488, 550, and 647 wavelengths (appears white in the overlay images) was considered to be background lipofuscin and was not analyzed. Aside from adjusting brightness/contrast, we performed no digital image processing to subtract background. We have previously attempted to optimize automated imaging analysis tools for our purposes, but these tools were designed to work with fresh, low background rodent tissues, not human samples taken from older organ donors. As such, we chose to implement a manual approach in our imaging analysis in which we used our own judgement of the negative/positive controls and target images to assess mRNA label.

For the RNAscope experiments, the same analysis procedure was conducted as previously described (8). 2-3 20X images were acquired of each human DRG section, and 3 sections were imaged per human donor. The raw image files were brightened and contrasted in Olympus CellSens software (v1.18), and then analyzed manually one neuron at a time for expression of each mRNA. **I**mages were not analyzed in a blinded fashion. Cell diameters were measured using the polyline tool. Total neuron counts were acquired by counting all probe-labeled neurons and all neurons that were clearly outlined by DAPI (satellite cell) signal and contained lipofuscin in the overlay image. For each section, the neuronal counts for each target/subpopulation from all images were summed, and the population percentages were calculated for that section. The population percentages from 3 sections were then averaged to yield the final population value for each donor. Pie-charts represent the average of all of three donors. The total number of neurons assessed is indicated in the Figure captions and represents the sum of all neurons analyzed between all 3 donors. Since *TRPV1* and *SCN10A* signal was assessed independently in three different experiments (in combination with *NTRK1*, *NTRK2*, and *NTRK3*), the data from all three experiments was combined. For those cases, if the same donor was used in each experiment, their population values were averaged. The same procedure applied to *CALCA* which was used multiple times (in combination with *PRDM12*, *SST* and *NPPB*).

For the RNAscope experiment shown in Figure 4 (*CALCA*/*NPPB* in males vs females), ∼10 *NPPB*-positive neurons from each section were imaged at 40X magnification, and 3 sections were imaged per donor (totaling ∼30 neurons/donor). The 40X images were then cropped to show only a single *NPPB*-positive neuron and the file names were blinded by a non-affiliated person. The blinded experimenter brightened and contrasted the images in Olympus CellSens and then drew regions of interest (ROIs) around the soma (not to include the larger mass of lipofuscin) (**Figure S8**). The area of the *CALCA* mRNA signal within the ROI was analyzed using the Count and Measure tool which highlights the mRNA puncta using a thresholded detection. Since RNAscope fluorescence intensity reflects the number of probe pairs bound to each molecule, a manual threshold was applied to each image so that all mRNA signal was highlighted within the ROI. Since each mRNA puncta in the ACD protocol averages 1.5µm^2^, the *CALCA* area measurements were divided by 1.5 to determine the number of puncta and then divided by the area of the ROI to yield *CALCA* mRNA puncta/µm^2^. Given the low-expression and detection of *NPPB* in the human DRG, we combined all the data from all donors (5 males, 3 females) which is reflected in the graph. Graphs were generated using GraphPad Prism version 8 (GraphPad Software, Inc. San Diego, CA USA). A relative frequency distribution histogram with a fitted Gaussian distribution curve was generated using the diameters of all mRNA-positive neurons detected in all experiments. Images in the Figures are pseudocolored.

### Human DRG cultures and electrophysiology

Human DRG cultures were prepared as described (83) from 3 donors (donors 2, 6, and 7). All electrophysiology experiments were performed between DIV5 and DIV7. Experiments were performed using a MultiClamp 700B (Molecular Devices) patch-clamp amplifier and PClamp 9 acquisition software (Molecular Devices) at room temperature. Recordings were sampled at 20 kHz and filtered at 3 kHz (Digidata 1550B, Molecular Devices). Pipettes (outer diameter, 1.5 mm; inner diameter, 1.1 mm, BF150-110-10, Sutter Instruments) were pulled using a PC-100 puller (Narishige) and heat polished to 2-3 MΩ resistance using a microforge (MF-83, Narishige). Series resistance was typically 5 MΩ and was compensated up to 70%. Data were analyzed using Clampfit 10 (Molecular Devices). All neurons included in the analysis had a resting membrane potential (RMP) more negative than −40 mV. In current-clamp mode, cells were held at RMP for the duration of the experiment. The pipette solution contained the following (in mM): 120 K-gluconate, 6 KCl, 4 ATP-Mg, 0.3 GTP-Na, 0.1 EGTA, 10 HEPES and 10 phosphocreatine, pH 7.2 (adjusted with N-methyl glucamine), and osmolarity was ∼290 mOsm. The external solution contained the following (in mM): 135 NaCl, 2 CaCl_2_, 1 MgCl_2_, 5 KCl, 10 glucose, and 10 HEPES, pH 7.4 (adjusted with N-methyl glucamine), and osmolarity was adjusted to ∼315 mOsm with sucrose. Cells were dialyzed for 3-5 minutes after break-in with the internal solution before commencing recordings. The cells were continuously perfused with the external solution using a ValveLink 8 perfusion system. Stock solution of capsaicin was diluted to 200 nM in the external solution and applied directly to the patched neuron using the perfusion system. Following 10 s of baseline recording of spontaneous activity in current clamp mode, capsaicin was applied for 10 s to test for depolarization of the neuron or action potential generation.

### Selection of neuronal barcodes in Loupe Browser

We manually selected all barcoded spots that overlapped neurons in Loupe Browser (v4.2.0, 10x Genomics) and exported as csv files for each sample. Surrounding barcodes were computationally obtained based on neuronal barcodes’ coordinates. To avoid duplicates and keep data consistent, barcodes could only have one classification. For instance, if a surrounding barcode was also overlapping a neuron, it was removed from the surrounding barcodes. By exclusion, barcodes that were not labeled neuronal or were not directly surrounding a barcode were labeled as ‘other barcodes’. For downstream analysis, we used neuronal barcodes that overlapped only single neurons. We observed that 3.19% of these neuronal barcodes overlapped the same neuron, with 0.66% being assigned to different clusters. After determination of neuronal clusters, all H&E images of each donor section was loaded into Loupe Browser and the barcodes for the identified neuronal clusters in that section were remapped for visualization purposes. An image of the overlaid barcodes on the tissue section was saved and then stacked with the high-resolution H&E image in CellSens. Each neuron and barcode was visualized on the Loupe Browser image, and then the same neuron was found by toggling to the high-resolution image. Neuronal diameters for neurons with visible nuclei were then measured using the polyline tool.

### Human and macaque transcriptome comparison

Based on comparative transcriptomic analysis of mouse and human DRG RNA-seq, we previously found that neuronal subtype-restricted genes were likely to be conserved in expression in the hDRG bulk RNA-seq data (5). Hence, as a starting point for analysis of conservation of lineage-restricted gene expression across subpopulations in human and macaque DRGs, we first identified the top 555 neuronal lineage-restricted coding genes in the hDRG (genes in the lowest 5% of normalized entropy signifying tissue restricted expression, **File S30**) out of 11,117 medium or high expression genes in the Visium dataset (read count >= 3 in one or more cells). Gene expression in the macaque orthologs in Smart-seq2 assay were obtained from literature (23), but many of these genes have low dynamic range (abundance between 0 – 0.1 across subpopulations) likely due to the nature of the Smart-seq assay. 111 lineage-restricted human DRG genes with higher dynamic range in macaque Smart-seq data were used to perform clustering of human and macaque subpopulations (based on expression enrichment scores in subpopulations for these genes), followed by clustering of the genes based on their gene expression patterns to assess conservation of lineage-restricted gene expression patterns in the two species. Of these, 91 genes in 3 modules are shown in Figure 6.

### Statistical Analysis

#### Visium spatial RNA-seq analysis

Raw count data for the selected neuronal barcodes was obtained from the respective feature-barcode matrices. We used Python (v3.7 with Anaconda distribution), R (v4.0.3) and Seurat (v3.2.2) for data analysis. Prior to initiating Seurat clustering workflow, data was cleaned by removing barcodes with low counts (<2000). We verified that selected neuronal barcodes that had no expression (count<1) of the neuronal marker *SNAP25,* had minimal overlap with neurons, and for that reason they were also excluded from downstream analysis. The remaining 3,952 neuronal barcodes, grouped by donor ID, created a total of eight Seurat objects. The standard Seurat integration workflow was followed (28). This integration workflow can reduce batch effects by identifying pairwise correspondences (named “anchors”) between single barcodes across samples. First, each object was normalized and identified the 2,000 most variable features. After identifying anchors, the data was integrated, generating one combined Seurat object. Next, the data was scaled, and the combined Seurat object was further processed following the standard Seurat clustering workflow. Clustering and visualization were performed using Uniform Manifold Approximation and Projection (UMAP) as the dimensionality reduction algorithm. After the first round of clustering, some clusters had these non-neuronal genes as cluster markers: *APOD, MT3, MPZ, CPLX1, SPARCL1, IFI27, CKB, BLVRB, SPP1, VSNL1, CLEC2L, CEND1, TECR, HSPB6, SNRNP25, SNCB, FAM57B, ATP1A3, NAT8L, MGP, TAGLN, DEXI, FABP7, TIMP1, CD74, VIM.* These genes were influencing the clustering and to overcome this, we scored the non-neuronal signal using ‘AddModuleScore’ function. This function scores the difference between the average expression levels of each gene set and randomly selected control gene set, across the neuronal barcodes (84). The barcodes were re-clustered (resolution=1) and the non-neuronal signal was regressed out (vars.to.regress=nn_score1). We identified markers for each cluster using the Wilcoxon Rank Sum test integrated in Seurat. Some pairs of clusters had a set of neuronal markers that were unique with respect to all the other clusters but were shared between the two of them. Therefore, clusters without clear distinct neuronal markers were merged to generate the final clusters (**Figure S3**).

#### Differential expression analysis

In order to identify neuron-specific sex-differences, we conducted differential expression analysis in neuronal barcodes. Because our neuronal barcodes may contain signal originating in the surrounding cells, we performed statistical analysis for the surrounding barcodes. **Figure S7A** shows the number of neuronal barcodes and surrounding barcodes used for statistical analysis. We combined barcode counts to generate a pseudo bulk sample for each neuronal cluster, respective surrounding barcodes and overall neuronal and overall surrounding barcodes. This approach ensures that statistical hypothesis testing is applicable to the tested population of barcodes, and not subject to sampling variance within the large number of individual barcodes in each population. Additionally, any effect of varying amounts of neuronal mRNA proportion across spots also get homogenized by such pooling. Genes with less than 10 reads, were excluded from each combined sample and removed from downstream analysis. Each dataset was then analyzed by using DESeq2 (85), which normalized the raw gene counts (gene counts are divided by barcode-specific normalization factors that are calculated based on the median ratio of gene counts relative to geometric mean per gene) and corrected for batch effect followed by testing for differential abundance. We performed differential expression analysis using the ‘DESeq’ function (this function performs differential expression analysis based on the negative binominal distribution and Wald statistics). Nominal p-values were corrected for multiple testing using the Benjamini-Hochberg (BH) method (86). In addition, we performed shrinkage of the Log2 Fold Change (LFC) estimates in order to generate more accurate LFC. We used the adaptive shrinkage estimator from the ‘ashr’ R package (87) and set the contrast to male vs female as the groups we wanted to compare. Genes were considered to be differentially expressed (DE) if FC ≥ 1.33 and adjusted p-value ≤ 0.05. Since mRNA profiles in each spot is an admixture of multiple cell types, we considered a gene to be specifically DE in neuronal barcodes if it was not DE in surrounding barcodes. Statistical hypothesis testing results for all tests can be found in **Files S2-23**. For each gene tested we report baseMean (mean of normalized counts), log2FoldChange (log2 fold change), lfcSE (standard error), pvalue (Wald test p-value) and padj (BH adjusted p-values). NA represents missing values.

## Supporting information

Zip of Supplemental files 1 - 30

## Supplementary Materials

This manuscript includes the following supplementary materials:

**Figure S1.** Visium spatial RNA-seq statistics and approach to selecting neuronal barcodes.

**Figure S2.** VISIUM tissue optimization.

**Figure S3.** Clustering RNA profiles of selected neuronal barcodes.

**Figure S4.** Pruritogen receptor enriched and C-LTMR clusters.

**Figure S5.** Size distribution of neurons in each cluster.

**Figure S6.** RNAscope for *LPAR3*, *PVALB*, *TRPM8* and *PENK* in the human dorsal root ganglia.

**Figure S7.** Analysis of sex-differences.

**Figure S8.** Abundance analysis of *CALCA* mRNA using RNAscope.

**Figure S9:** Expression of interleukin and receptor genes in human and mouse datasets.

**Figure S10:** Expression of ASIC genes in human and mouse datasets.

**Figure S11:** Expression of anoctamin genes in human and mouse datasets.

**Figure S12:** Expression of aquaporins genes in human and mouse datasets.

**Figure S13:** Expression of calcium channel genes in human and mouse datasets.

**Figure S14:** Expression of chloride channel genes in human and mouse datasets.

**Figure S15:** Expression of cholinergic receptor genes in human and mouse datasets.

**Figure S16:** Expression of GABA receptor genes in human and mouse datasets.

**Figure S17:** Expression of gap-junction genes in human and mouse datasets.

**Figure S18:** Expression of glutamate receptor genes in human and mouse datasets.

**Figure S19:** Expression of glycine receptor genes in human and mouse datasets.

**Figure S20:** Expression of neuropeptide genes in human and mouse datasets.

**Figure S21:** Expression of potassium channel genes in human and mouse datasets.

**Figure S22:** Expression of purinergic receptor genes in human and mouse datasets.

**Figure S23:** Expression of transient receptor potential genes in human and mouse datasets.

**Figure S24:** Expression of neuronal transcription factors in human and mouse datasets.

**Figure S25:** Expression of understudied GPCRs in human and mouse datasets.

**Figure S26:** Expression of understudied ion channels in human and mouse datasets.

**Figure S27:** Expression of understudied kinases in human and mouse datasets.

**Figure S28:** Expression of low entropy genes.

**Table S1.** Human donor information.

**Table S2.** Summary of RNAscope experiment details.

**File S1. (separate Excel file)**

Ranked gene expression within each neuronal barcode by cluster.

**Files S2-27. (separate Excel files)** Results of statistical analysis for sex differences.

**File S28. (separate Excel file)** Gene enrichment analysis for the genes DE in pruritogen receptor enriched population.

**File S29. (separate Excel file)** Overall sequencing statistics for Visium experiments.

**File S30. (separate Excel file)** Data underlying human and macaque DRG gene expression comparison.

## Acknowledgements

The authors thank the organ donors and their families for their enduring gift. We thank Erin Vines and the staff at the Southwest Transplant Alliance for coordinating DRG recovery from organ donation surgeries. We also thank members of the Price Lab, Moeno Kume, Dr. Muhammad Saad Yousuf, Dr. Amelia Balmain, Juliet Mwirigi for assistance in DRG recovery. The authors thank the Genome Center at The University of Texas at Dallas for the services to support our research. We thank Marcos Chavez and Jardel Kuate Fotso for creating the website for public access and visualization of our processed data.

## Funding

This work was supported by NIH grants NS111929 to PMD and TJP, NS065926 to GD and TJP and NS042595 to RWG IV.

## Author contributions

DT-F and SS conducted Visium spatial optimization and gene expression protocol. DT-F performed Visium spatial RNA-seq analysis, differential expression analysis and human Visium versus mouse single-cell comparison. PRR performed human and macaque transcriptome comparison and assisted with computational analyses. SS performed RNAscope and analysis. VJ performed electrophysiology. AW assisted in computational analyses. IS assisted in Visium experiments. AC, JCR and BC trained staff on DRG surgical extraction and established DRG recovery protocols. AC and JCR did DRG recovery surgery. DT-F, SS and TJP wrote the manuscript. DT-F, SS, PRR, PMD, RWG IV, MDB, GD and TJP designed the study. PRR, GD and TJP supervised the study. PMD, RWG IV, GD and TJP obtained funding for the study. All authors read and edited the paper.

## Declaration of Interests

PRR, AW, GD and TJP are founders of Doloromics. The authors declare no other conflicts of interest.

## Data and materials availability

Raw sequencing data will be deposited in the dbGaP repository. Public access to processed data is available at sensoryomics.com. All analyses scripts and loupe browser files are available upon request.

## Ethics approval

This work was approved by the University of Texas at Dallas Institutional Review Board. Original approval date for protocol MR 15-237 -*The Human Dorsal Root Ganglion Transcriptome* was Sept 21, 2015. The protocol was renewed and has a current expiration date of Jan 22, 2023.

## Notes

### Summary of Updates

This version of the manuscript has been revised to add identification of additional sensory neuron clusters in our datasets, add additional analysis on drug target expression and add comparison to non-human primate DRG which was published after we posted the original version of the manuscript.

