## Supplemental Figures File for "Spatial transcriptomics reveals unique molecular fingerprints of human nociceptors"

Theodore J Price PhD

University of Texas at Dallas

800 W Campbell Rd

Richardson, TX 75080

972-883-4311

**This PDF file includes:**

Figures S1-S28

Tables S1 and S2

Captions for files S1 to S30

**Other Supplementary Materials for this manuscript include the following:**

Files S1-S30 (separate Excel files)

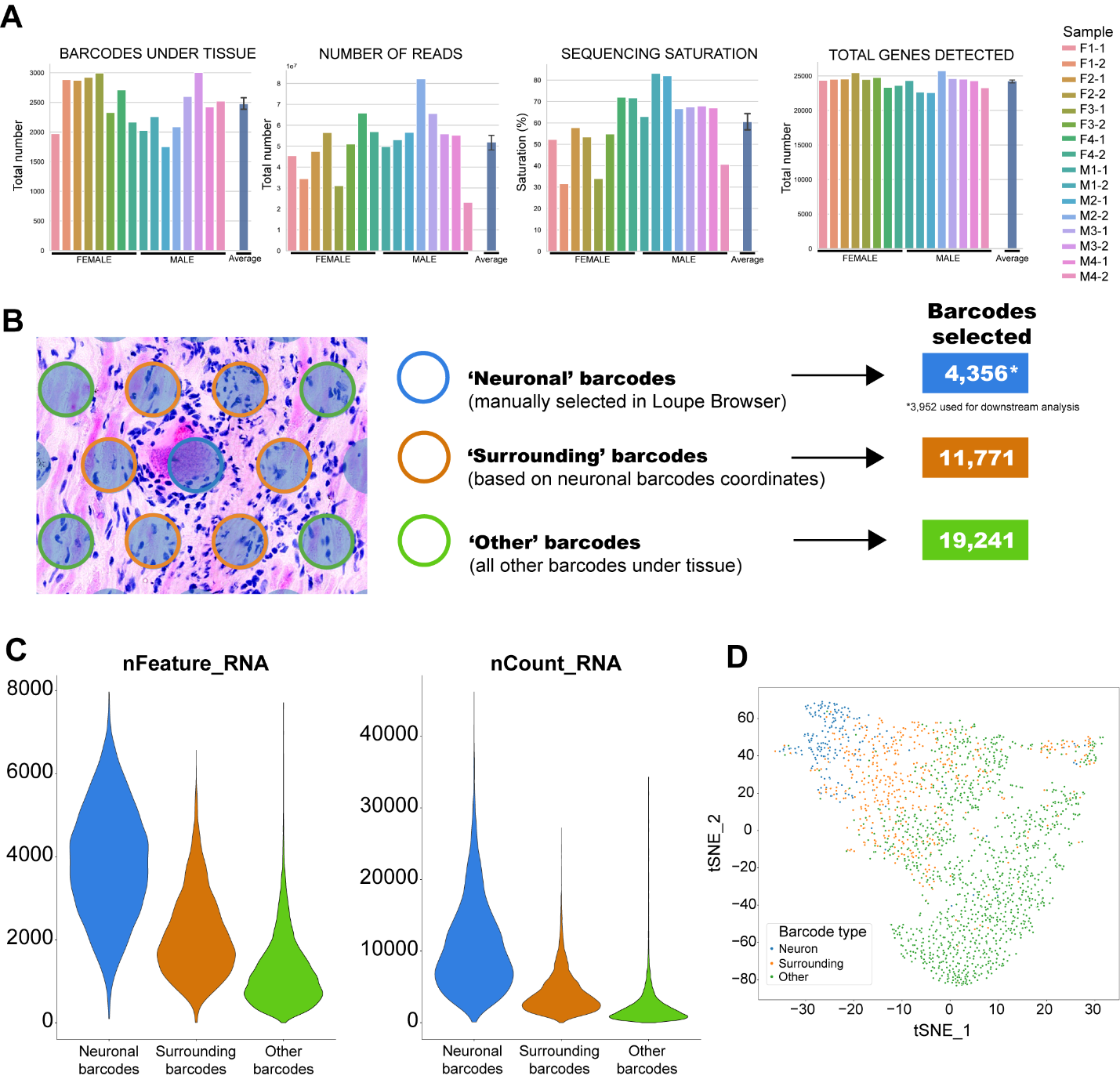

**Figure S1: Visium spatial RNA-seq statistics and approach to selecting neuronal barcodes**. **(A)** Summary of sequencing statistics for Visium spatial RNA-seq. Detailed statistics can be found in **File S25**. Sequencing saturation refers to the fraction of library complexity sequenced in each experiment, as estimated by Space Ranger. **(B)** Approach used to select neuronal barcodes for downstream analysis. Barcodes overlapping a single neuron were selected in Loupe Browser and barcodes directly surrounding each neuronal barcode were generated based on the neuronal barcodes’ coordinates. Remaining barcodes were labelled ‘other barcodes’. See Methods for details on barcode selection and classification. **(C)** Violin plots show that neuronal barcodes have higher number of detected genes (nFeature) and higher number of unique RNA molecules (nCount) than surrounding and other barcodes. **(D)** Representative tSNE (t-distributed Stochastic Neighbor Embedding) plot showing that neuronal barcodes have distinct profile compared to surrounding and other barcodes.

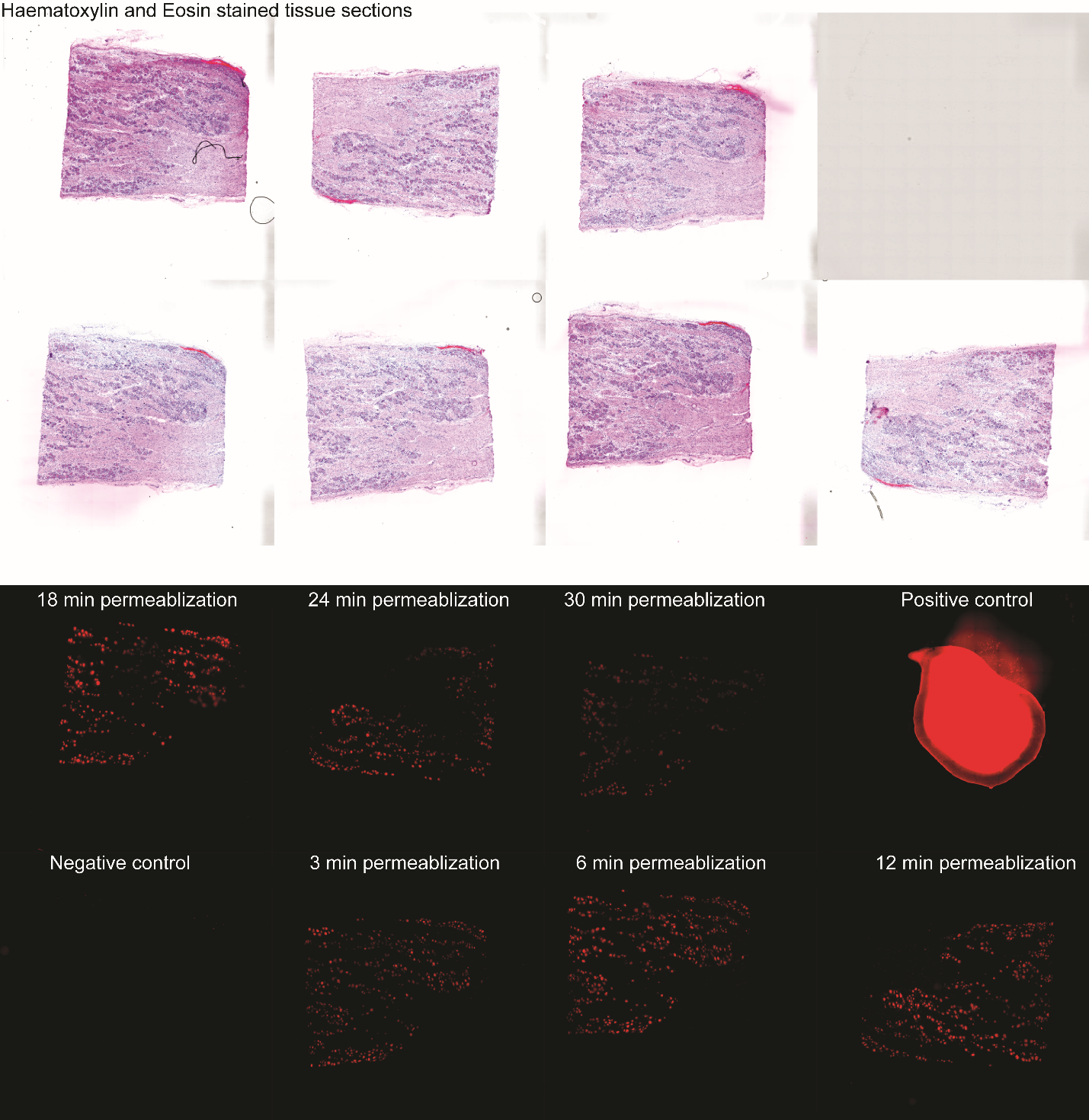

**Figure S2: VISIUM tissue optimization.** We conducted VISIUM spatial transcriptomics optimization protocol as instructed by 10X Genomics. In brief, sections of human DRG were placed onto etched frames on an optimization slide. The tissues were stained with Haematoxylin and Eosin and imaged (top panel). Tissues were then permeabilized and the mRNA was visualized using fluorescence microscopy (bottom panel). The positive control frame contained a drop of total mouse RNA. We detected bright and intense mRNA signal in neurons using the 6, 12, and 18 minute permeabilization times. As such, we used a 12-minute permeabilization for our spatial gene expression experiment.

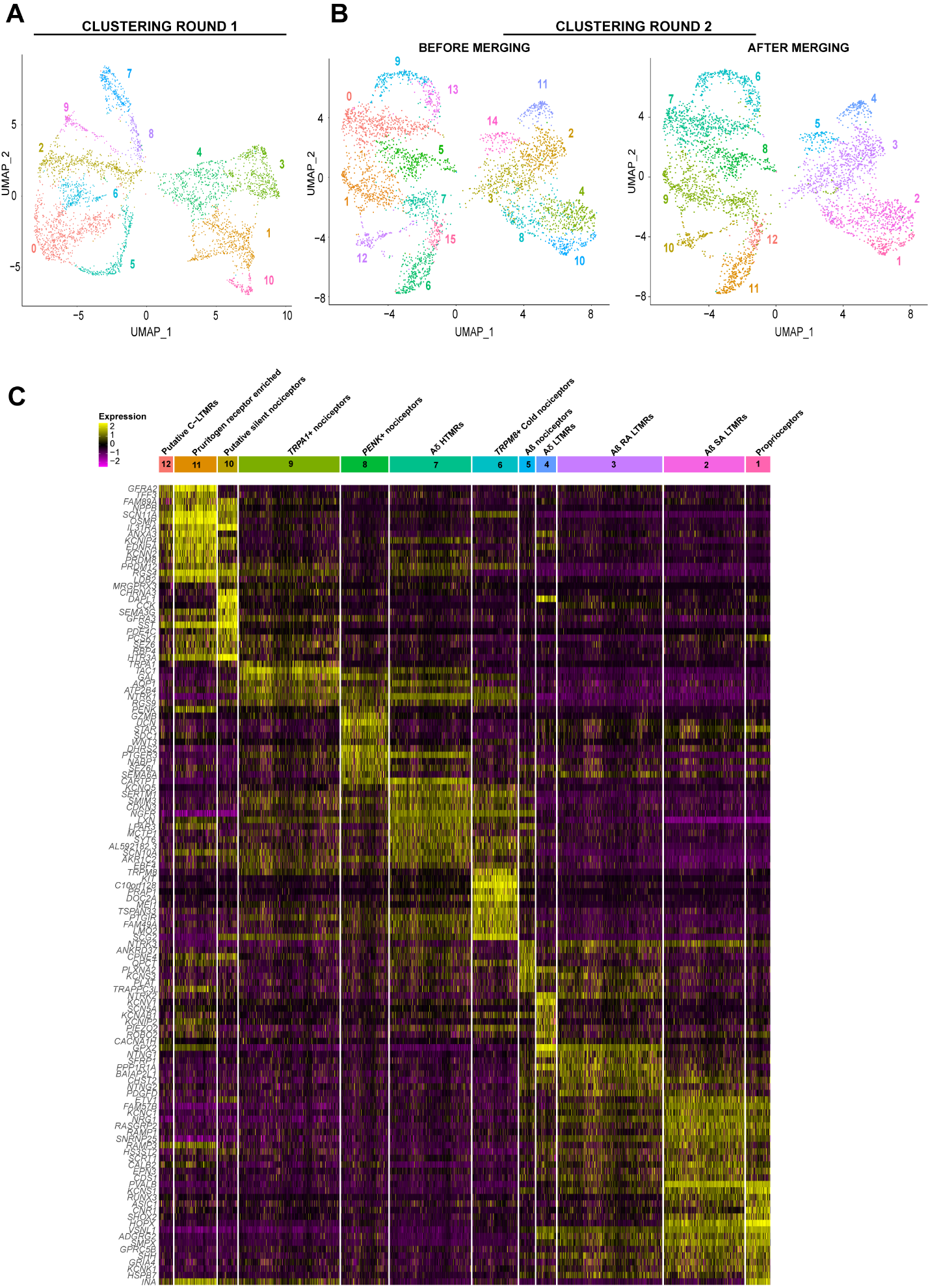

**Figure S3: Clustering RNA profiles of selected neuronal barcodes**. **(A)** UMAP plot shows first round of clustering that generated 11 clusters. Because of non-neuronal signal potentially coming from surrounding cells, non-neuronal genes that were enriched in all clusters were scored and regressed out in the next round of clustering. **(B)** Second round of clustering produced 16 clusters (B-left UMAP plot). Cluster without any distinguishable markers were merged, producing the final 12 clusters (B-right UMAP plot). (C) Heatmap showing expression of top neuronal marker genes in each cluster.

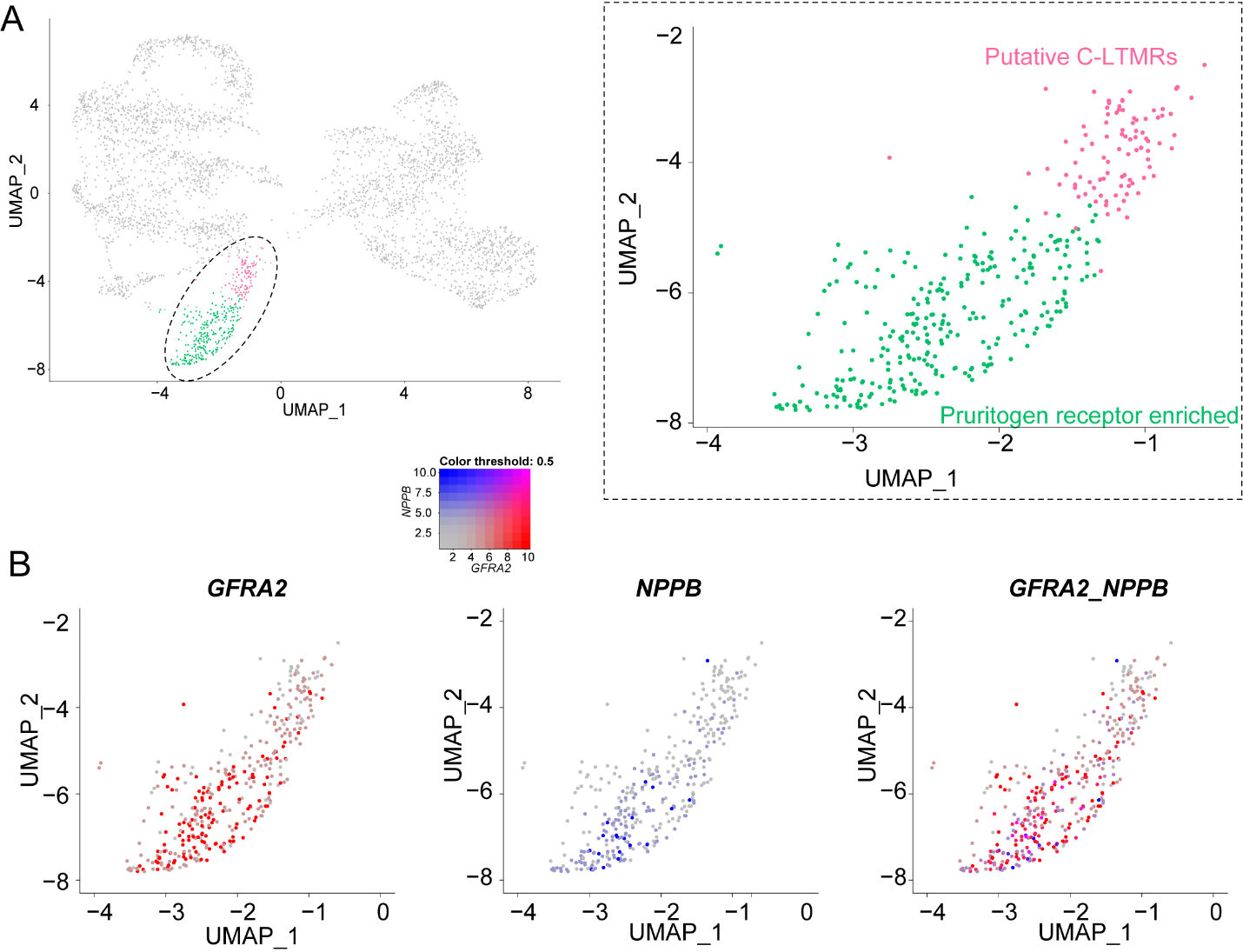

**Figure S4: Pruritogen receptor enriched and C-LTMR clusters. (A)** UMAP plots show two clusters that were generated in the second round of clustering. **(B)** Expression of *GFRA2* and *NPPB* across the two clusters. Expression of both genes was scaled from 1-10.

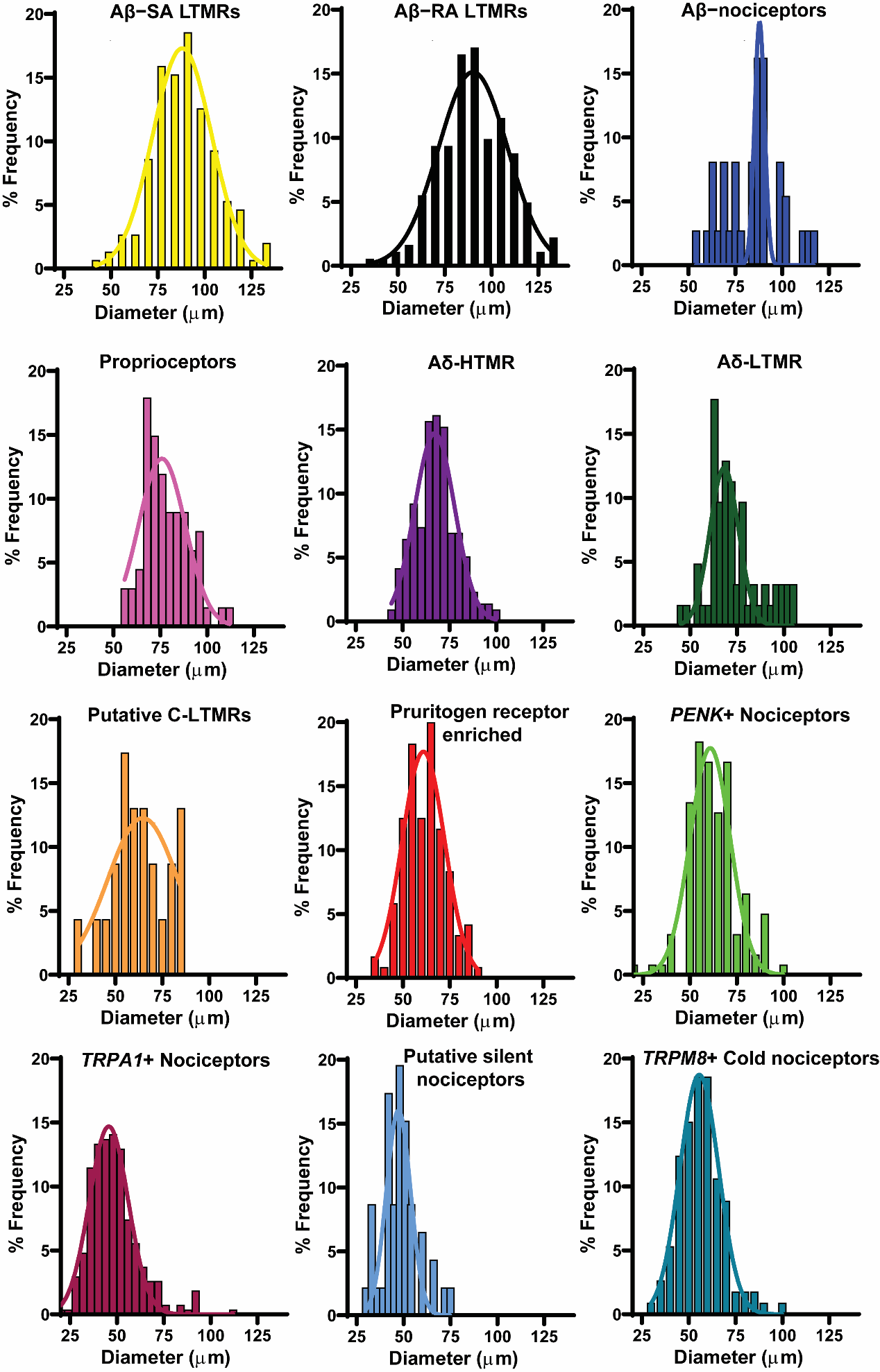

**Figure S5: Size distribution of neurons in each cluster.** The barcodes defining each cluster were remapped onto three DRG sections and the diameters of those neurons were measured. Gaussian fitted curve is also shown.

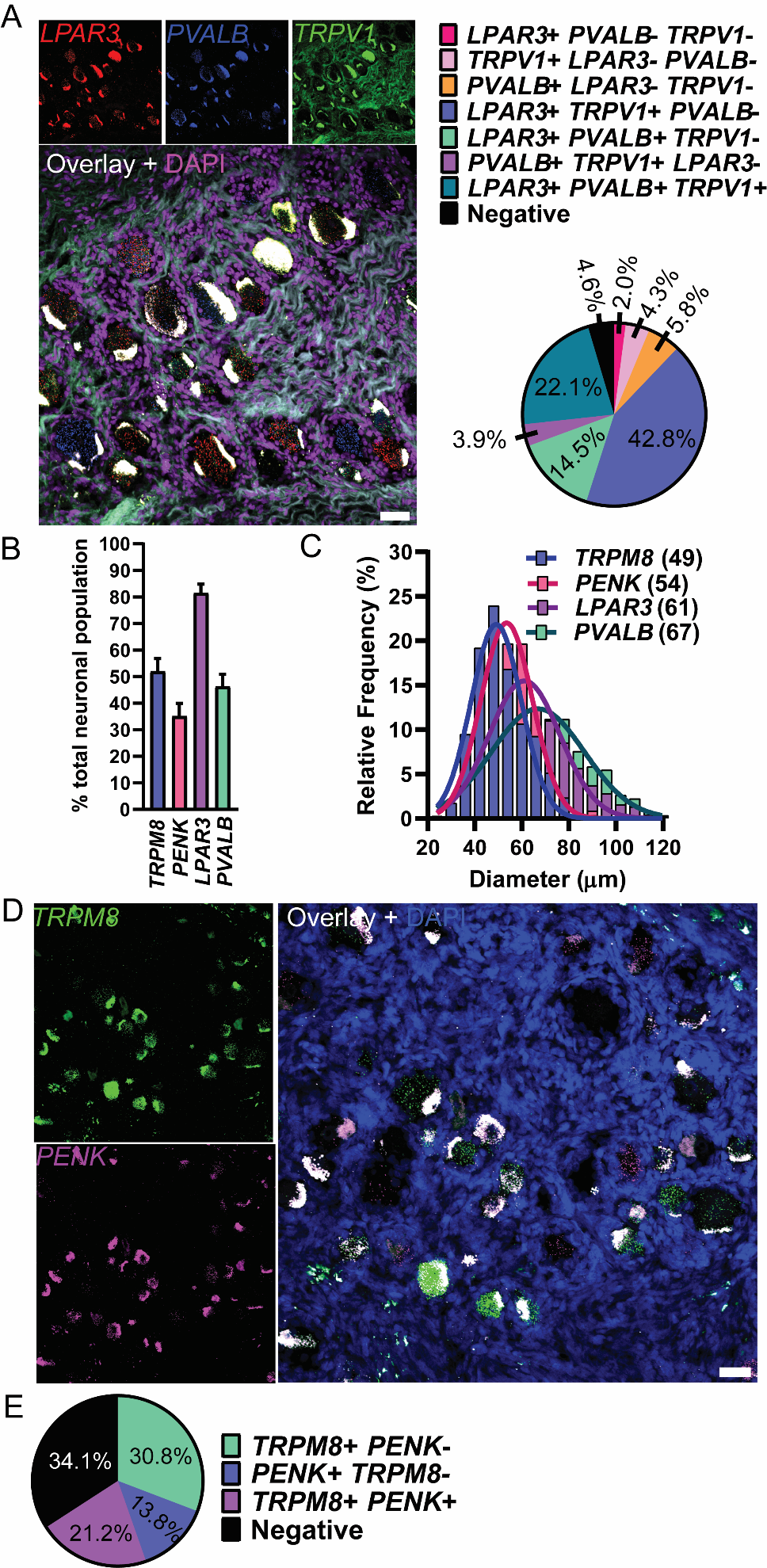

**Figure S6: RNAscope for *LPAR3, PVALB, TRPM8* and *PENK* in the human dorsal root ganglia. (A)** Representative image of *LPAR3*, *PVALB*, and *TRPV1* mRNA expression in the human DRG. Population distribution of each neuronal marker is shown in the pie chart. **(B)** Percentage of sensory neurons expressing *TRPM8*, *PENK*, *LPAR3*, and *PVALB* using RNAscope *in situ* hybridization. **(C)** The size distribution of all *TRPM8*, *PENK*, *LPAR3*, and *PVALB*-positive neurons. **(D)** Representative image of *TRPM8* and *PENK* mRNA expression in the human DRG. **(E)** Population distribution of *TRPM8* and *PENK* is shown in the pie chart. Scale bar = 50µm.

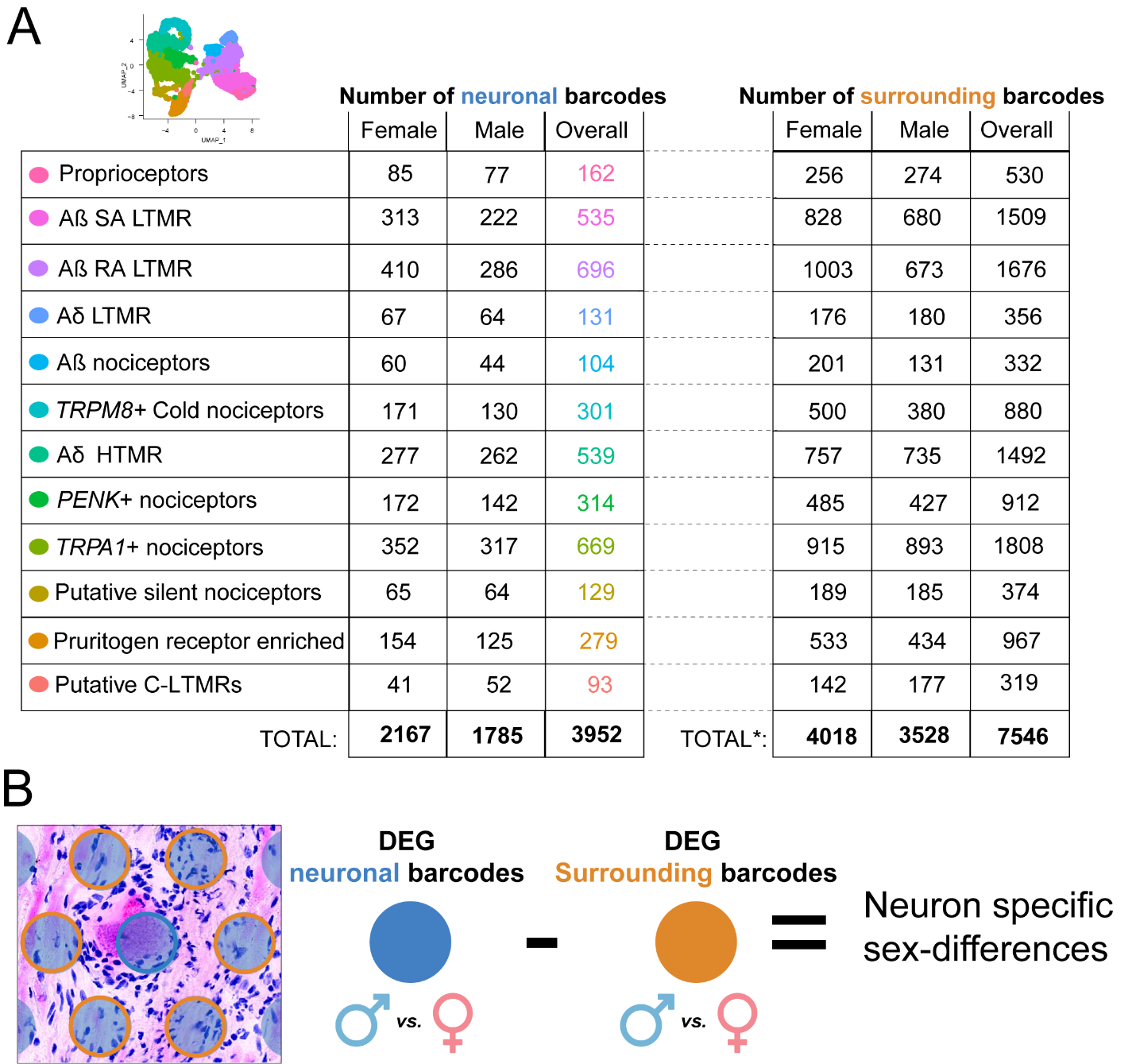

**Figure S7: Analysis of sex-differences. (A)** Number of barcodes used per cluster and overall, for neuronal barcodes and respective surrounding barcodes. *Note that the same barcodes may surround different neuronal clusters and, thus, the total number of surrounding barcodes used per respective cluster do not sum up to the total number of surrounding barcodes used for the overall analysis (duplicate barcodes were excluded). **(B)** Illustration of how the barcodes were identified from the spatial registered barcodes. DEG – differentially expressed genes.

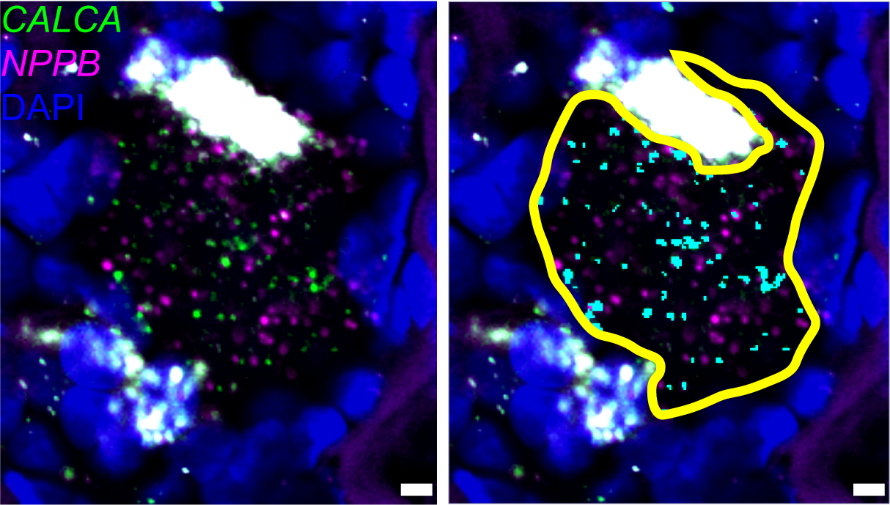

**Figure S8. Abundance analysis of *CALCA* mRNA using RNAscope.** On the left is a representative image of a single *NPPB*-positive (magenta) neuron co-labeled for *CALCA* mRNA (green) and DAPI (blue). As shown in the right image, a region of interest (yellow) was drawn around the neuronal cell body excluding the larger lipofuscin mass (white). *CALCA* signal was highlighted using a manual threshold in the Count and Measure tool in Olympus CellSens (cyan). The output of the tool is the sum of the highlighted area.

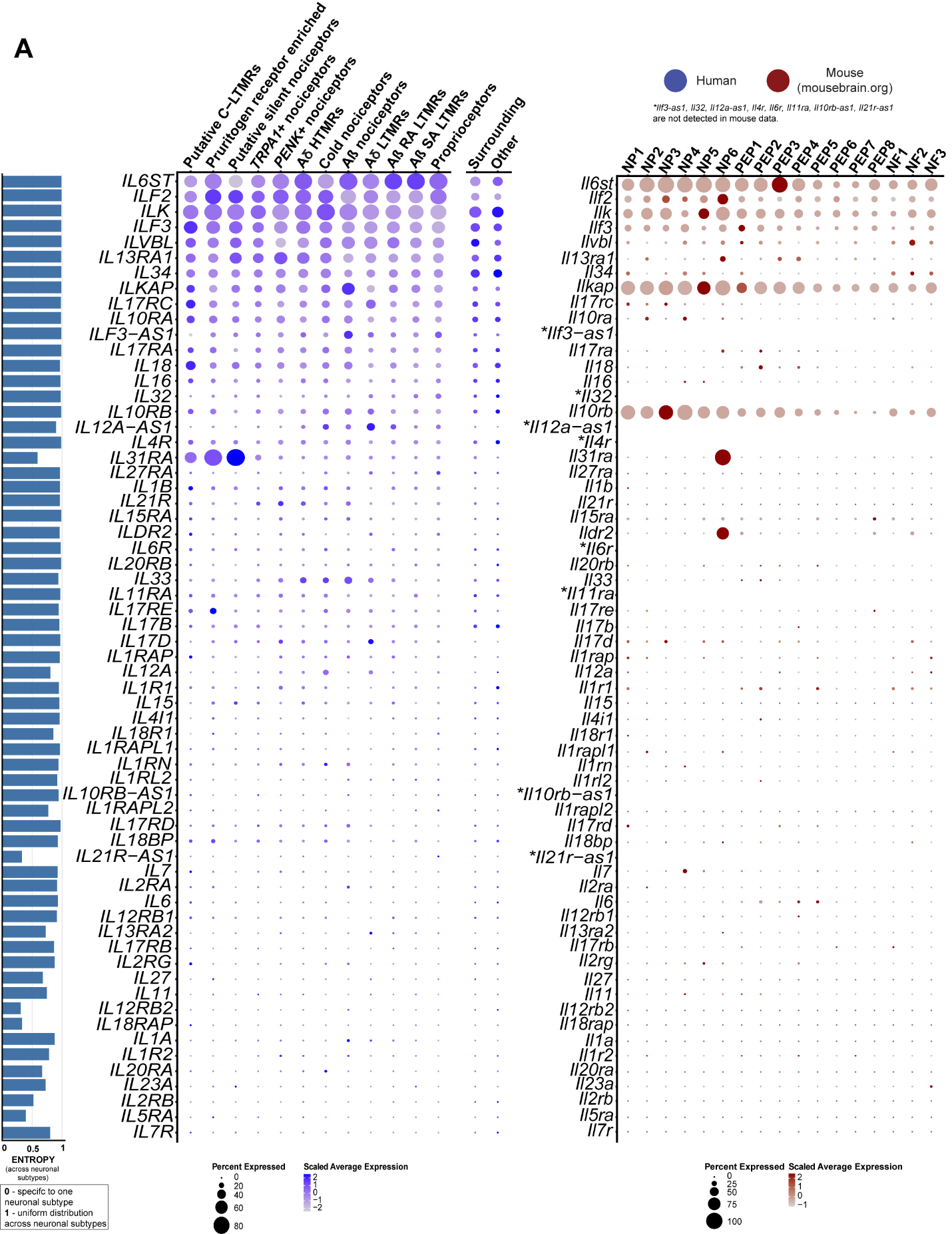

**Figure S9: Expression of** **interleukin and receptor genes in human and mouse datasets. (A)** Dot-plots showing the expression of interleukin and receptor genes in human spatial transcriptomic (in blue) and mouse single-cell experiments (in red). The size of the dot represents the percentage of barcodes within a cluster and the color corresponds to the average expression (scaled data) across all barcodes within a cluster for each gene shown. Normalized entropy was used as a measure of “specificity of neuronal subtype”, where a score of 0 means a gene is specific to one neuronal subtype and 1 means that a gene has uniform distribution across neuronal subtypes.

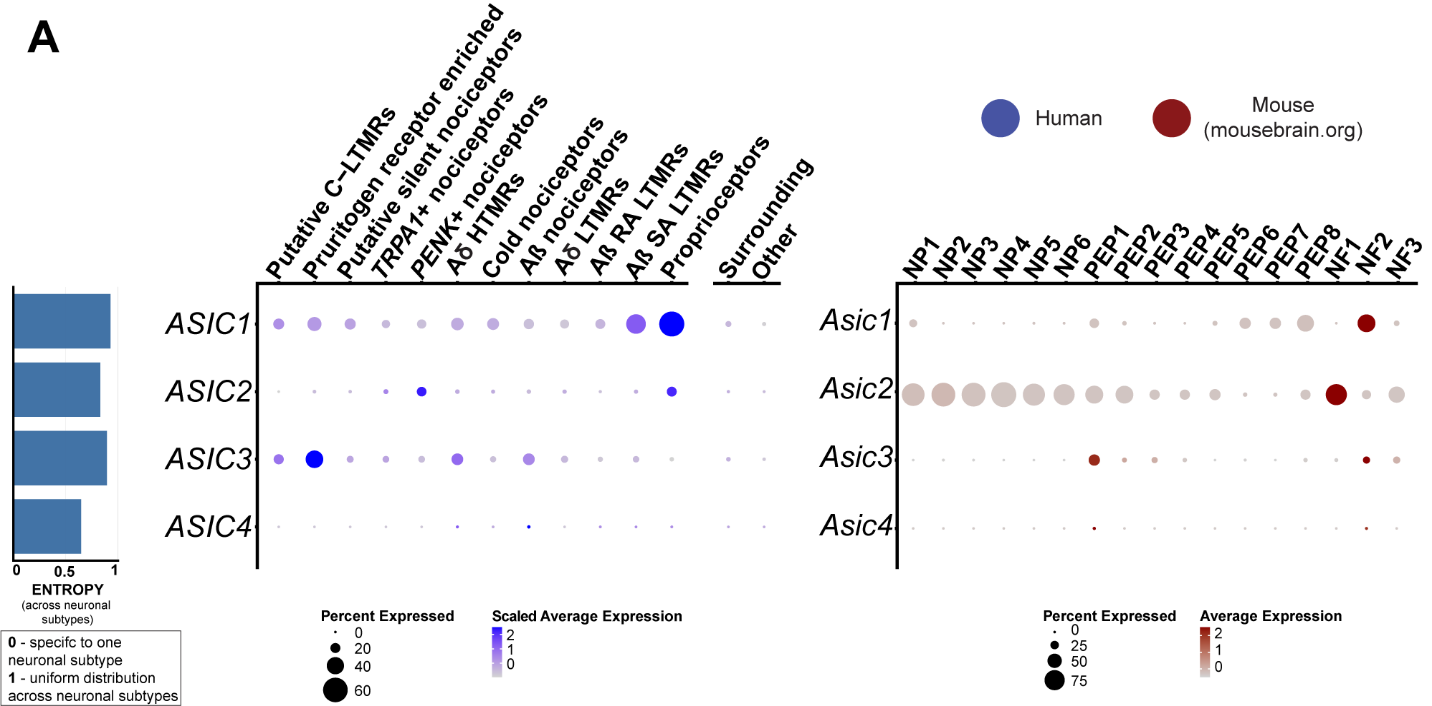

**Figure S10: Expression of ASIC genes in human and mouse datasets. (A)** Dot-plots showing the expression of ASIC genes in human spatial transcriptomic (in blue) and mouse single-cell experiments (in red). The size of the dot represents the percentage of barcodes within a cluster and the color corresponds to the average expression (scaled data) across all barcodes within a cluster for each gene shown. Normalized entropy was used as a measure of “specificity of neuronal subtype”, where a score of 0 means a gene is specific to one neuronal subtype and 1 means that a gene has uniform distribution across neuronal subtypes.

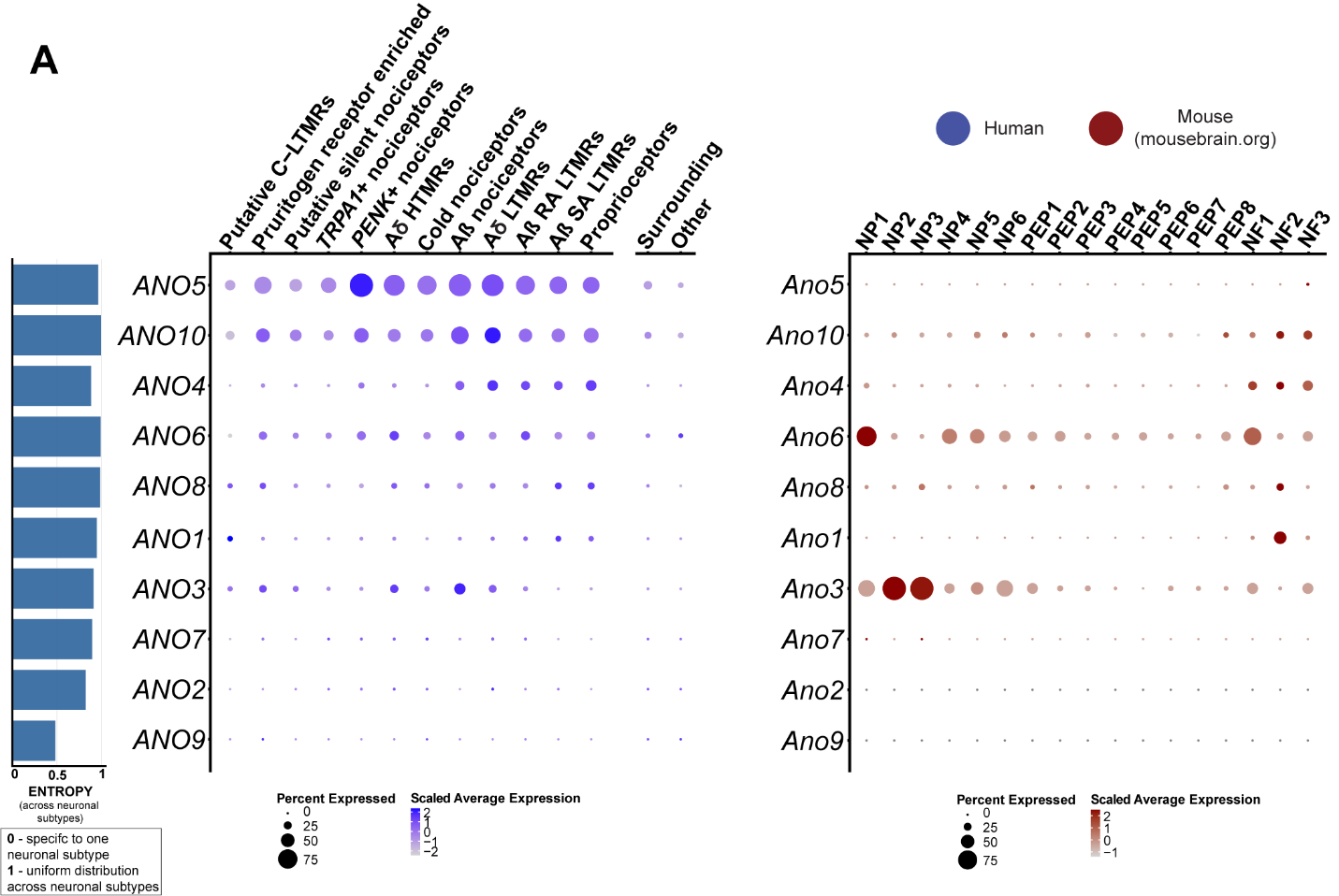

**Figure S11: Expression of anoctamin genes in human and mouse datasets. (A)** Dot-plots showing the expression of anoctamin genes in human spatial transcriptomic (in blue) and mouse single-cell experiments (in red). The size of the dot represents the percentage of barcodes within a cluster and the color corresponds to the average expression (scaled data) across all barcodes within a cluster for each gene shown. Normalized entropy was used as a measure of “specificity of neuronal subtype”, where a score of 0 means a gene is specific to one neuronal subtype and 1 means that a gene has uniform distribution across neuronal subtypes.

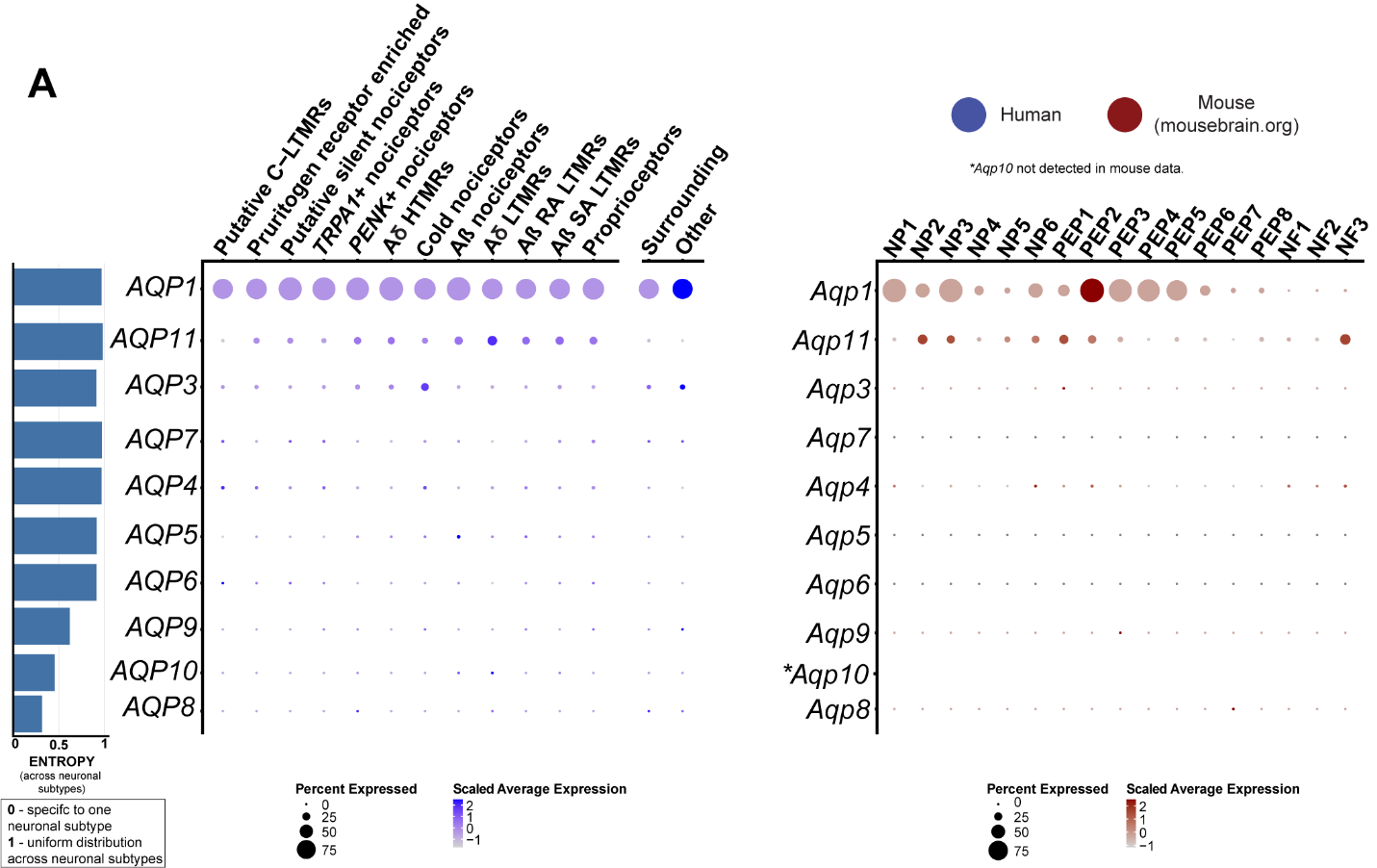

**F****igure S12: Expression of aquaporins genes in human and mouse datasets. (A)** Dot-plots showing the expression of aquaporins genes in human spatial transcriptomic (in blue) and mouse single-cell experiments (in red). The size of the dot represents the percentage of barcodes within a cluster and the color corresponds to the average expression (scaled data) across all barcodes within a cluster for each gene shown. Normalized entropy was used as a measure of “specificity of neuronal subtype”, where a score of 0 means a gene is specific to one neuronal subtype and 1 means that a gene has uniform distribution across neuronal subtypes.

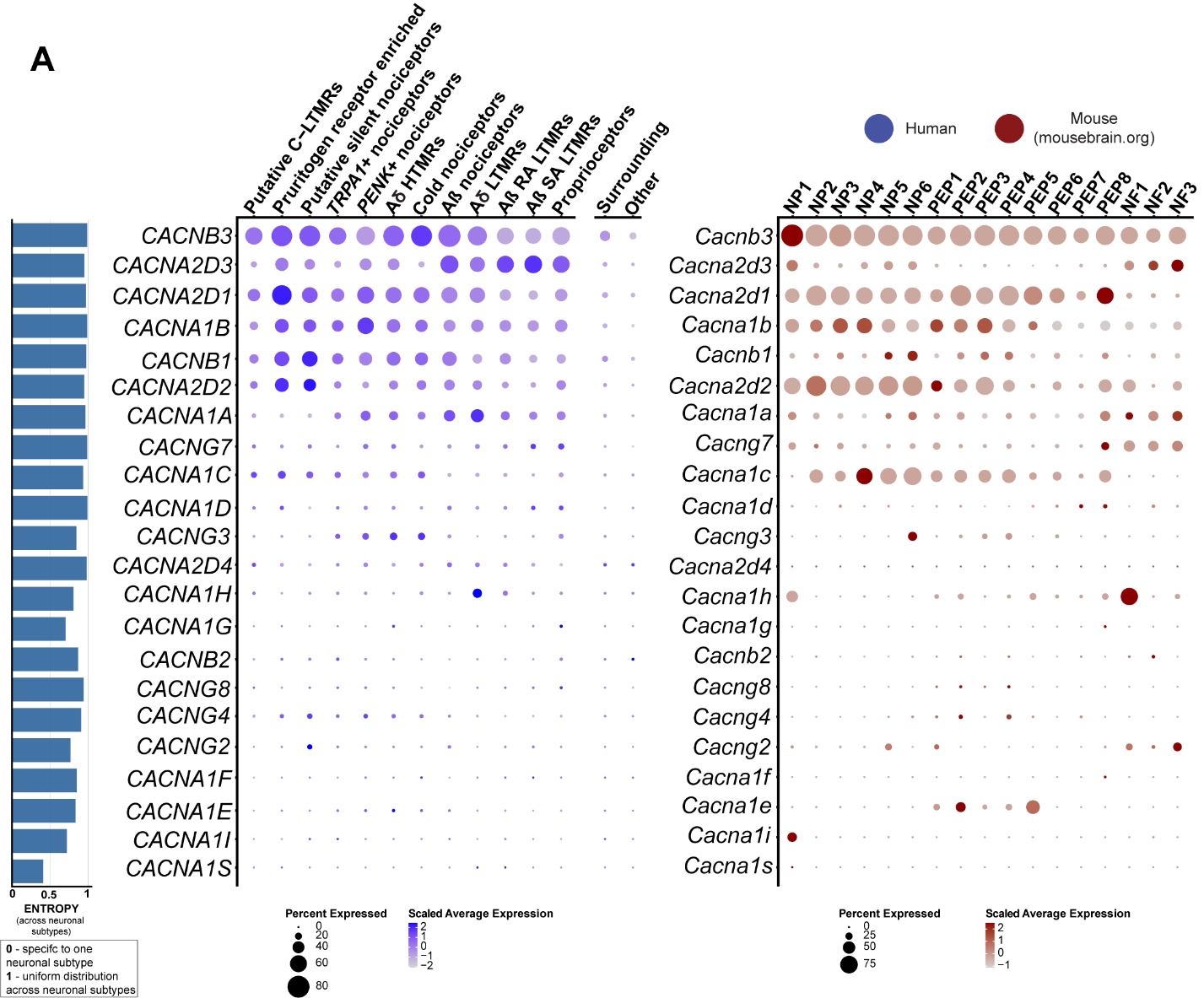

**Figure S13: Expression of calcium channel genes in human and mouse datasets. (A)** Dot-plots showing the expression of calcium channel genes in human spatial transcriptomic (in blue) and mouse single-cell experiments (in red). The size of the dot represents the percentage of barcodes within a cluster and the color corresponds to the average expression (scaled data) across all barcodes within a cluster for each gene shown. Normalized entropy was used as a measure of “specificity of neuronal subtype”, where a score of 0 means a gene is specific to one neuronal subtype and 1 means that a gene has uniform distribution across neuronal subtypes.

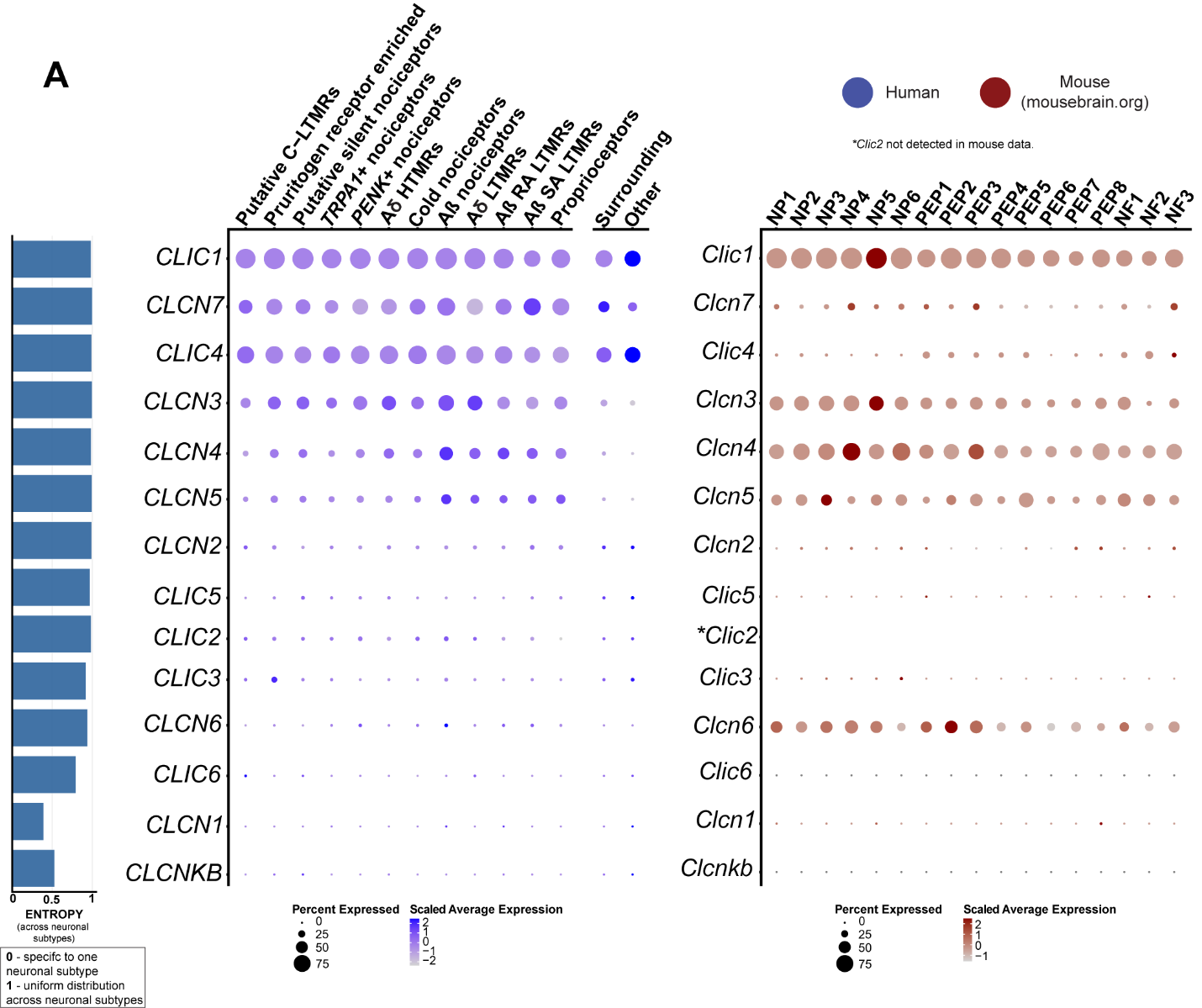

**Figure S14: Expression of chloride channel genes in human and mouse datasets. (A)** Dot-plots showing the expression of chloride channel genes in human spatial transcriptomic (in blue) and mouse single-cell experiments (in red). The size of the dot represents the percentage of barcodes within a cluster and the color corresponds to the average expression (scaled data) across all barcodes within a cluster for each gene shown. Normalized entropy was used as a measure of “specificity of neuronal subtype”, where a score of 0 means a gene is specific to one neuronal subtype and 1 means that a gene has uniform distribution across neuronal subtypes.

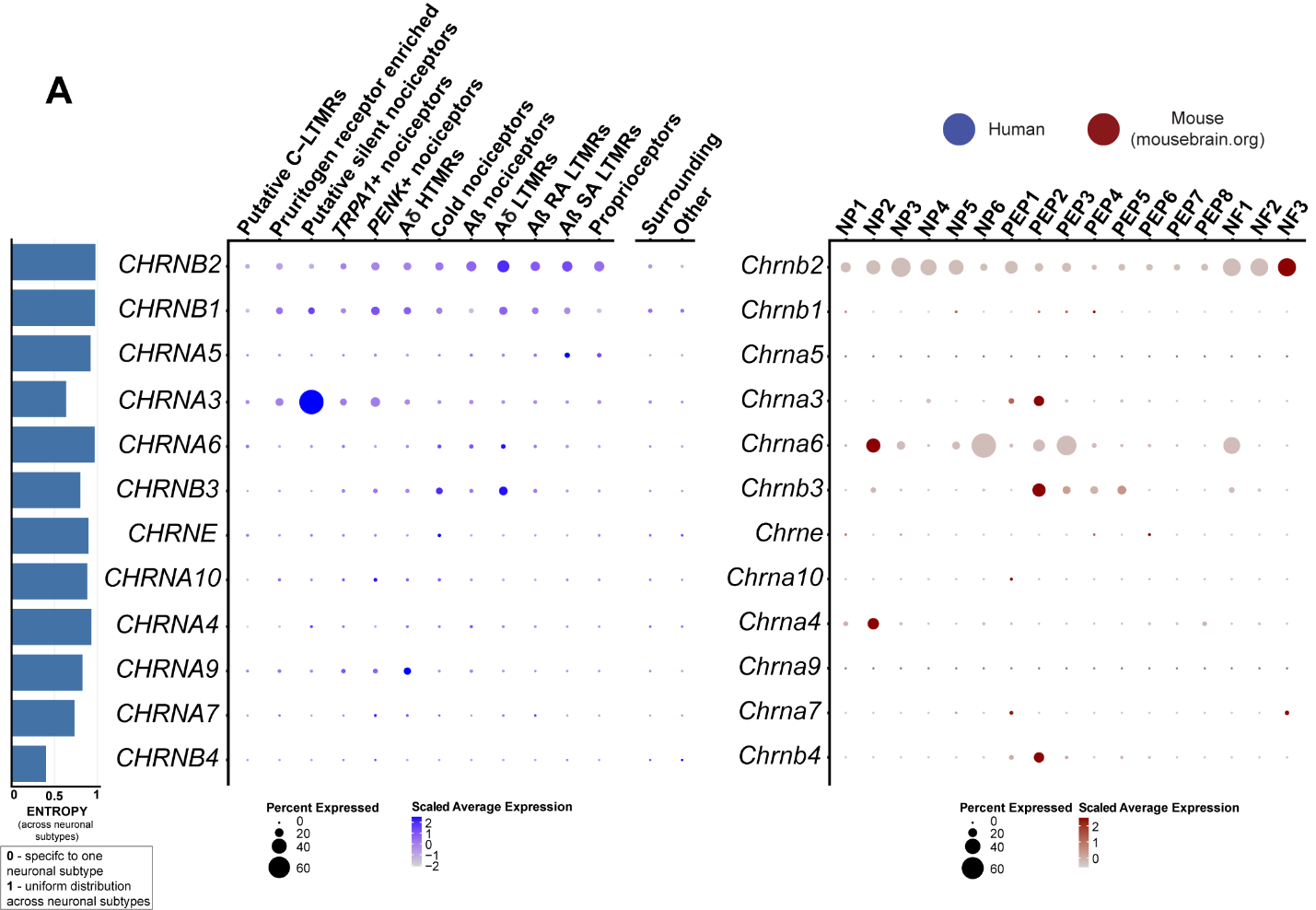

**Figure S15: Expression of cholinergic receptor genes in human and mouse datasets. (A)** Dot-plots showing the expression of cholinergic receptor genes in human spatial transcriptomic (in blue) and mouse single-cell experiments (in red). The size of the dot represents the percentage of barcodes within a cluster and the color corresponds to the average expression (scaled data) across all barcodes within a cluster for each gene shown. Normalized entropy was used as a measure of “specificity of neuronal subtype”, where a score of 0 means a gene is specific to one neuronal subtype and 1 means that a gene has uniform distribution across neuronal subtypes.

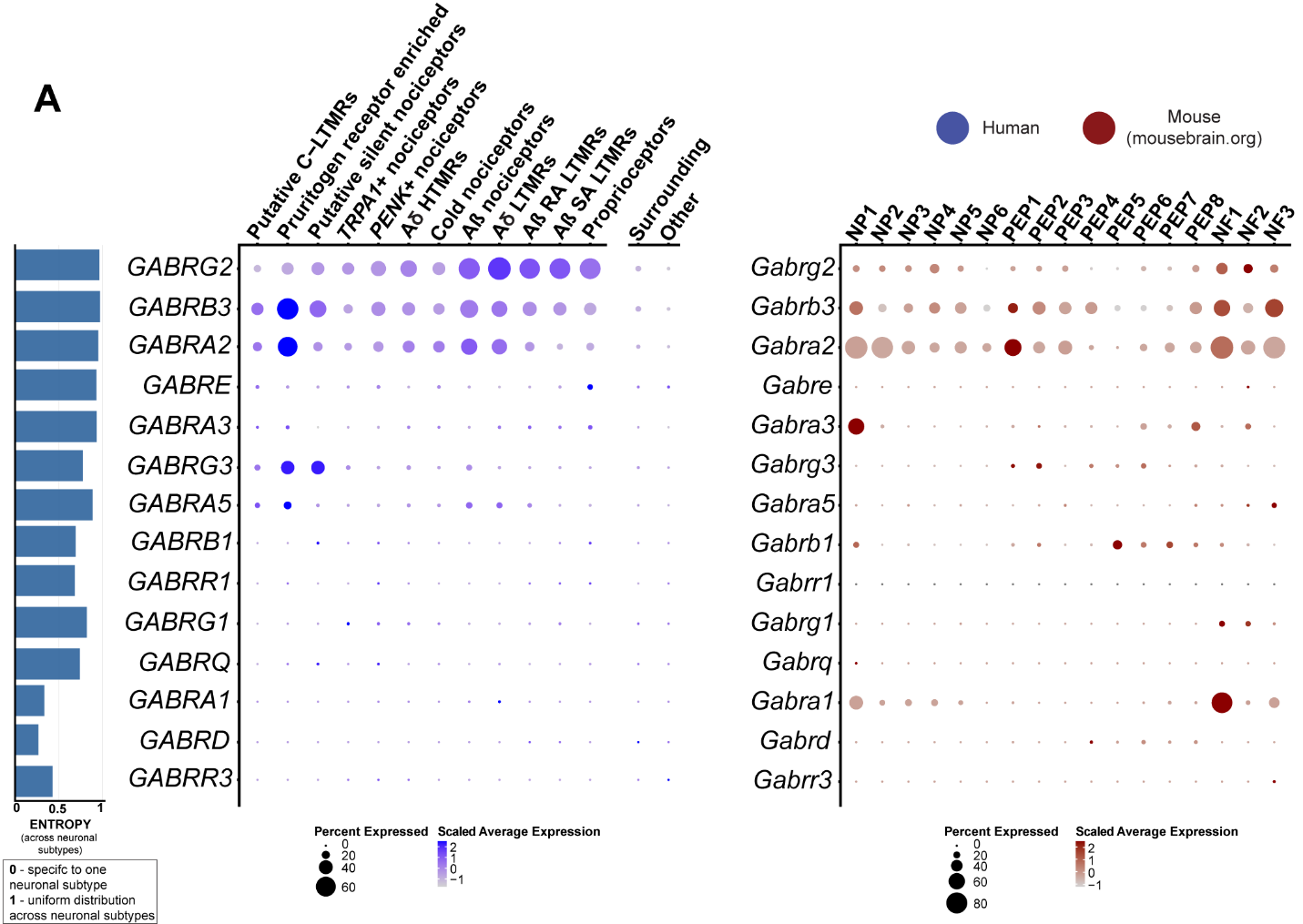

**Figure S16: Expression of ionotropic GABA receptor genes in human and mouse datasets. (A)** Dot-plots showing the expression of GABA receptor genes in human spatial transcriptomic (in blue) and mouse single-cell experiments (in red). The size of the dot represents the percentage of barcodes within a cluster and the color corresponds to the average expression (scaled data) across all barcodes within a cluster for each gene shown. Normalized entropy was used as a measure of “specificity of neuronal subtype”, where a score of 0 means a gene is specific to one neuronal subtype and 1 means that a gene has uniform distribution across neuronal subtypes.

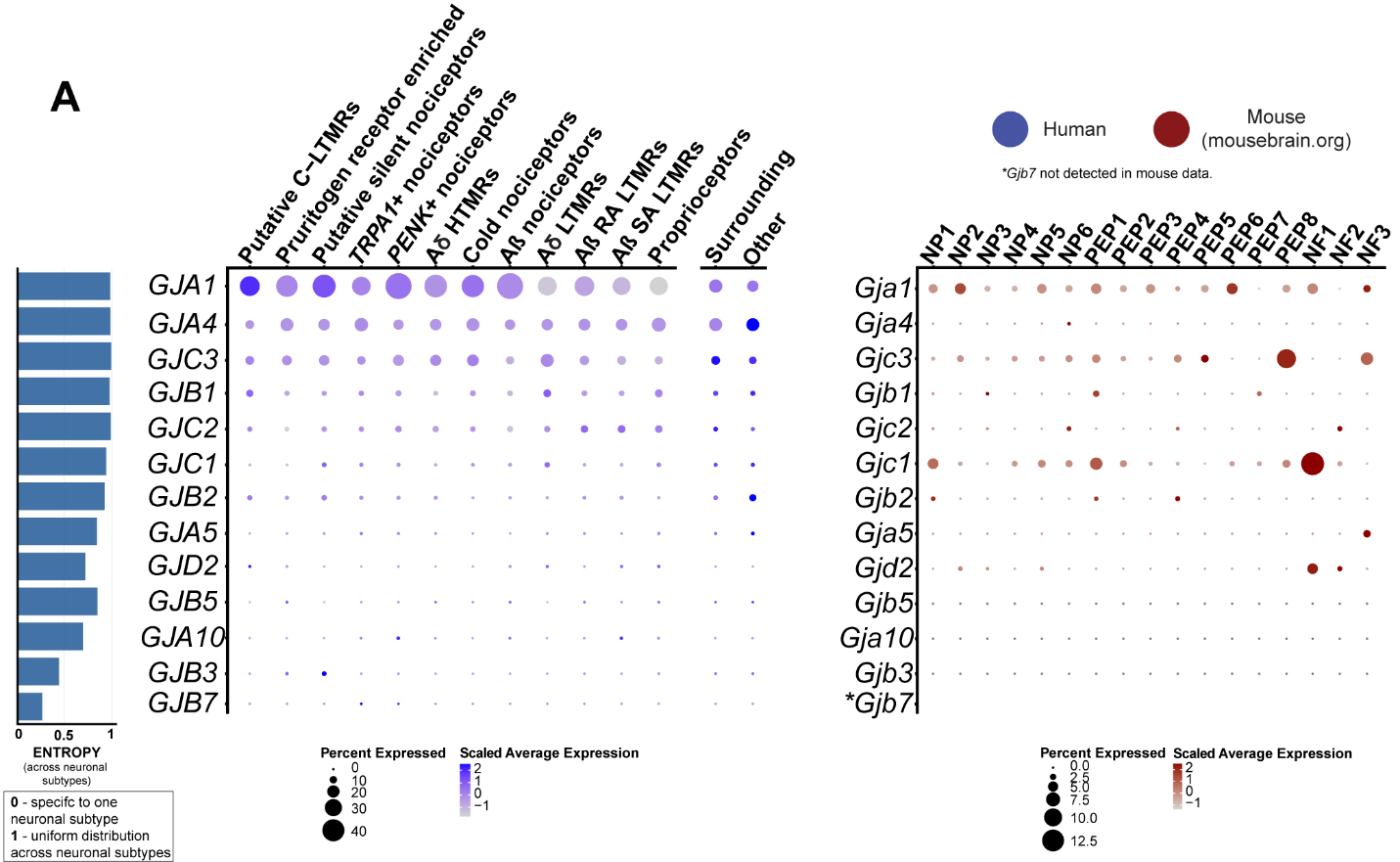

**Figure S17: Expression of gap-junction (connexin) genes in human and mouse datasets. (A)** Dot-plots showing the expression of gap-junction genes in human spatial transcriptomic (in blue) and mouse single-cell experiments (in red). The size of the dot represents the percentage of barcodes within a cluster and the color corresponds to the average expression (scaled data) across all barcodes within a cluster for each gene shown. Normalized entropy was used as a measure of “specificity of neuronal subtype”, where a score of 0 means a gene is specific to one neuronal subtype and 1 means that a gene has uniform distribution across neuronal subtypes.

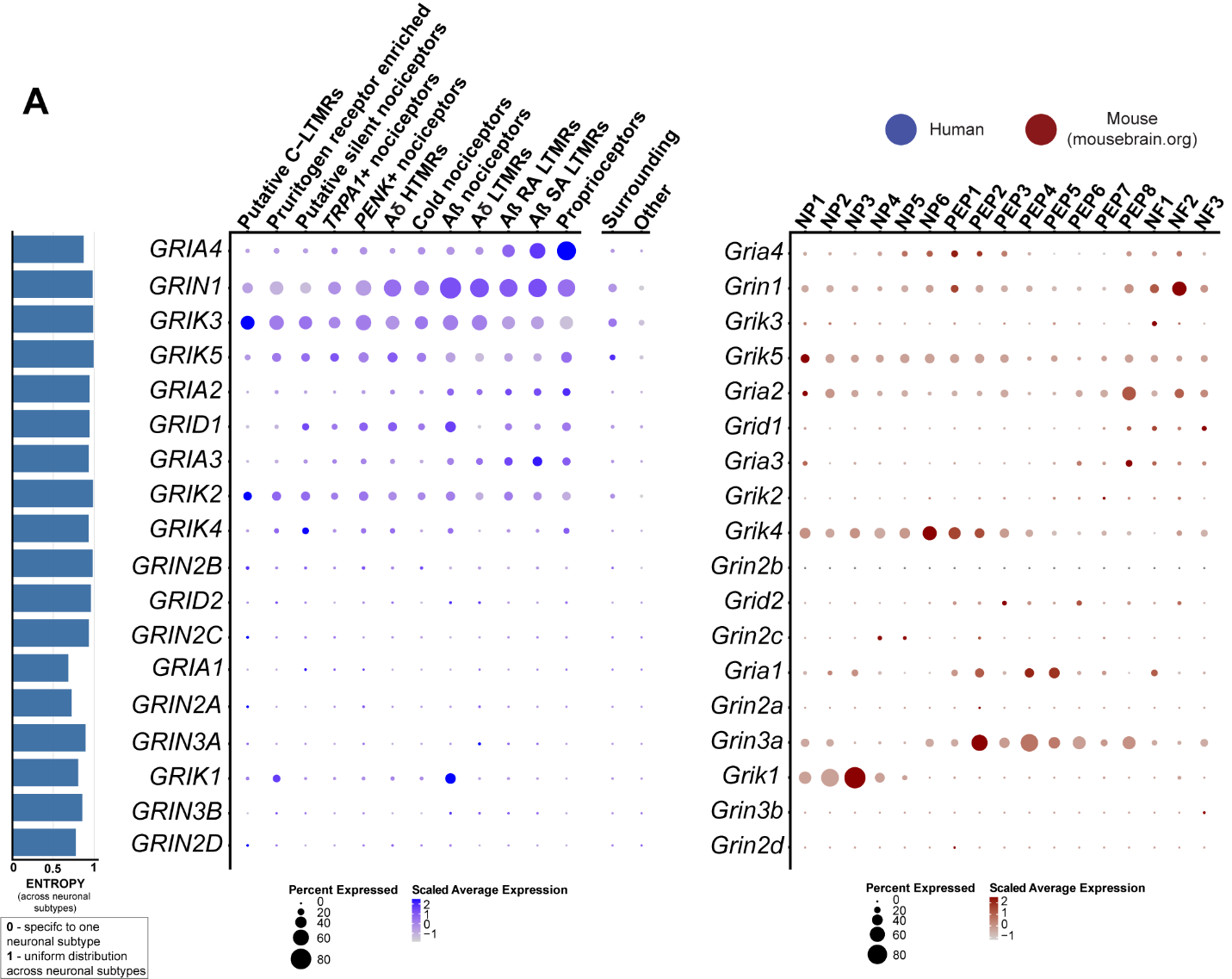

**Figure S18: Expression of ionotropic glutamate receptor genes in human and mouse datasets. (A)** Dot-plots showing the expression of glutamate receptor genes in human spatial transcriptomic (in blue) and mouse single-cell experiments (in red). The size of the dot represents the percentage of barcodes within a cluster and the color corresponds to the average expression (scaled data) across all barcodes within a cluster for each gene shown. Normalized entropy was used as a measure of “specificity of neuronal subtype”, where a score of 0 means a gene is specific to one neuronal subtype and 1 means that a gene has uniform distribution across neuronal subtypes.

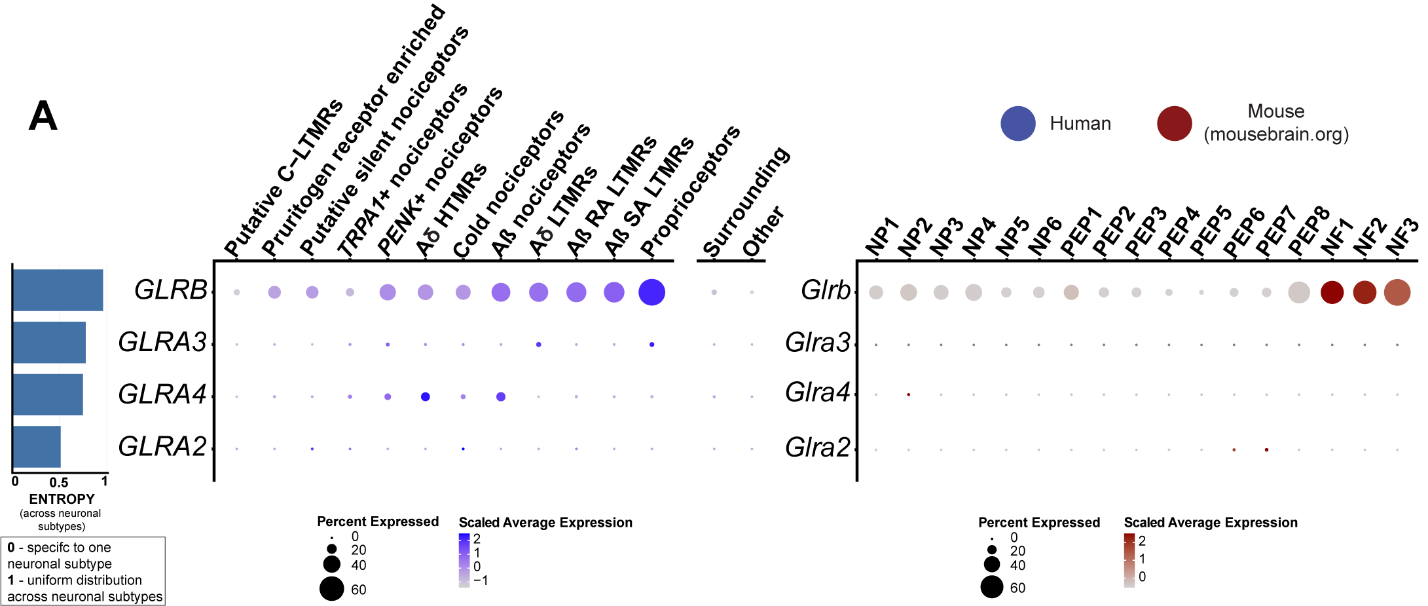

**Figure S19: Expression of glycine receptor genes in human and mouse datasets. (A)** Dot-plots showing the expression of glycine receptor genes in human spatial transcriptomic (in blue) and mouse single-cell experiments (in red). The size of the dot represents the percentage of barcodes within a cluster and the color corresponds to the average expression (scaled data) across all barcodes within a cluster for each gene shown. Normalized entropy was used as a measure of “specificity of neuronal subtype”, where a score of 0 means a gene is specific to one neuronal subtype and 1 means that a gene has uniform distribution across neuronal subtypes.

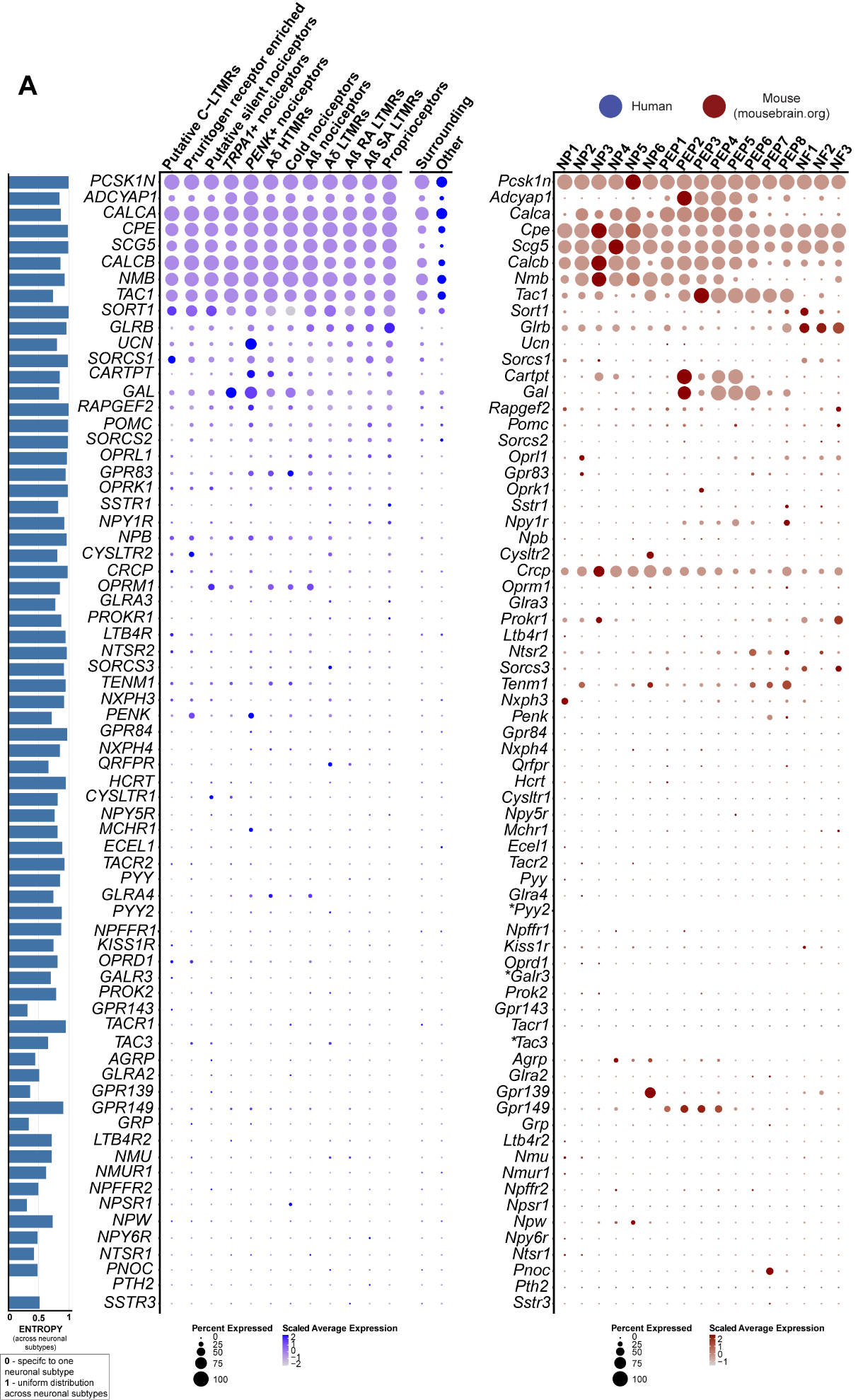

**Figure S20: Expression of neuropeptide genes in human and mouse datasets. (A)** Dot-plots showing the expression of neuropeptide genes in human spatial transcriptomic (in blue) and mouse single-cell experiments (in red). The size of the dot represents the percentage of barcodes within a cluster and the color corresponds to the average expression (scaled data) across all barcodes within a cluster for each gene shown. Normalized entropy was used as a measure of “specificity of neuronal subtype”, where a score of 0 means a gene is specific to one neuronal subtype and 1 means that a gene has uniform distribution across neuronal subtypes.

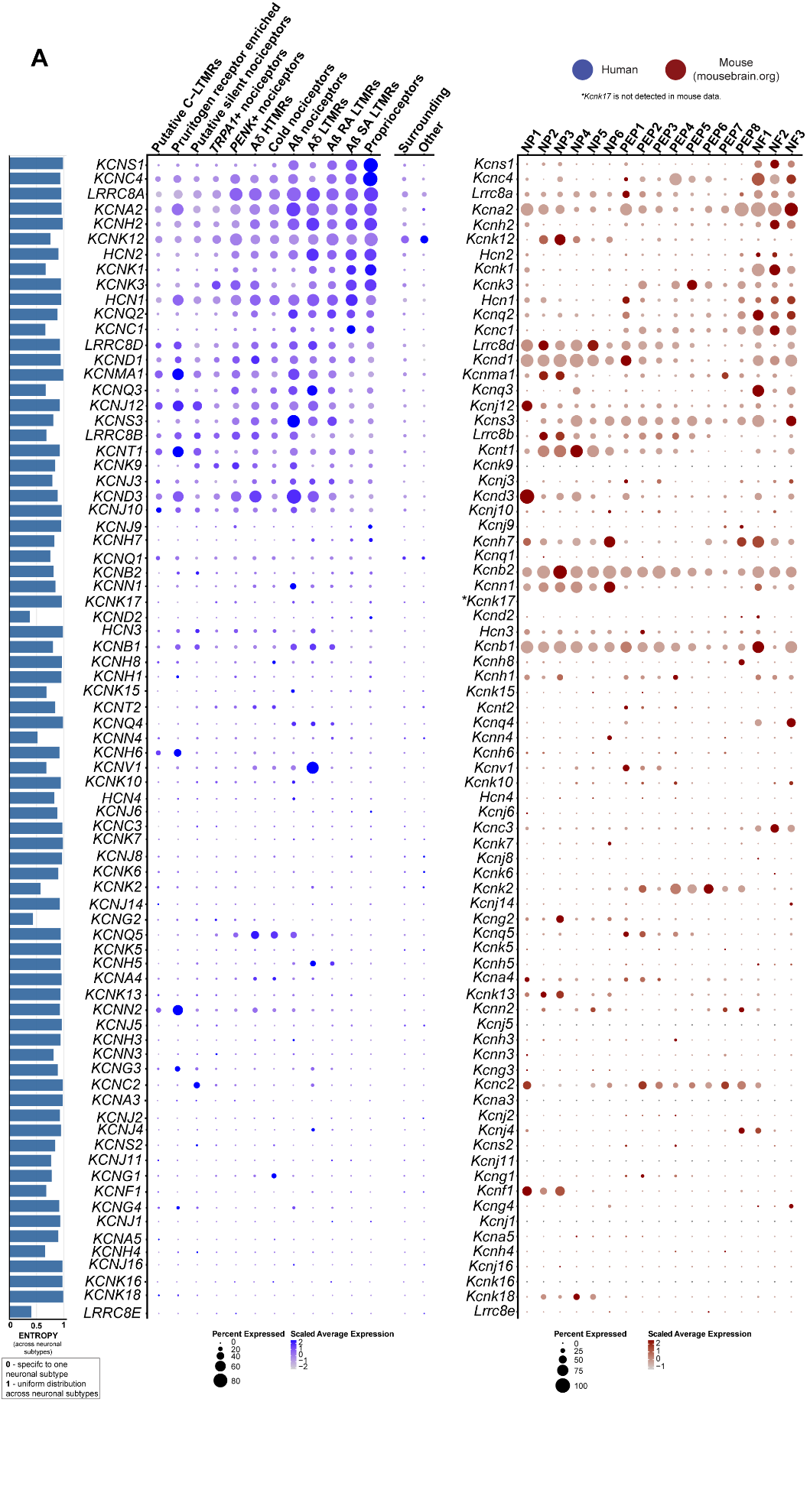

**Figure S21: Expression of potassium channel genes in human and mouse datasets. (A)** Dot-plots showing the expression of potassium channel genes in human spatial transcriptomic (in blue) and mouse single-cell experiments (in red). The size of the dot represents the percentage of barcodes within a cluster and the color corresponds to the average expression (scaled data) across all barcodes within a cluster for each gene shown. Normalized entropy was used as a measure of “specificity of neuronal subtype”, where a score of 0 means a gene is specific to one neuronal subtype and 1 means that a gene has uniform distribution across neuronal subtypes.

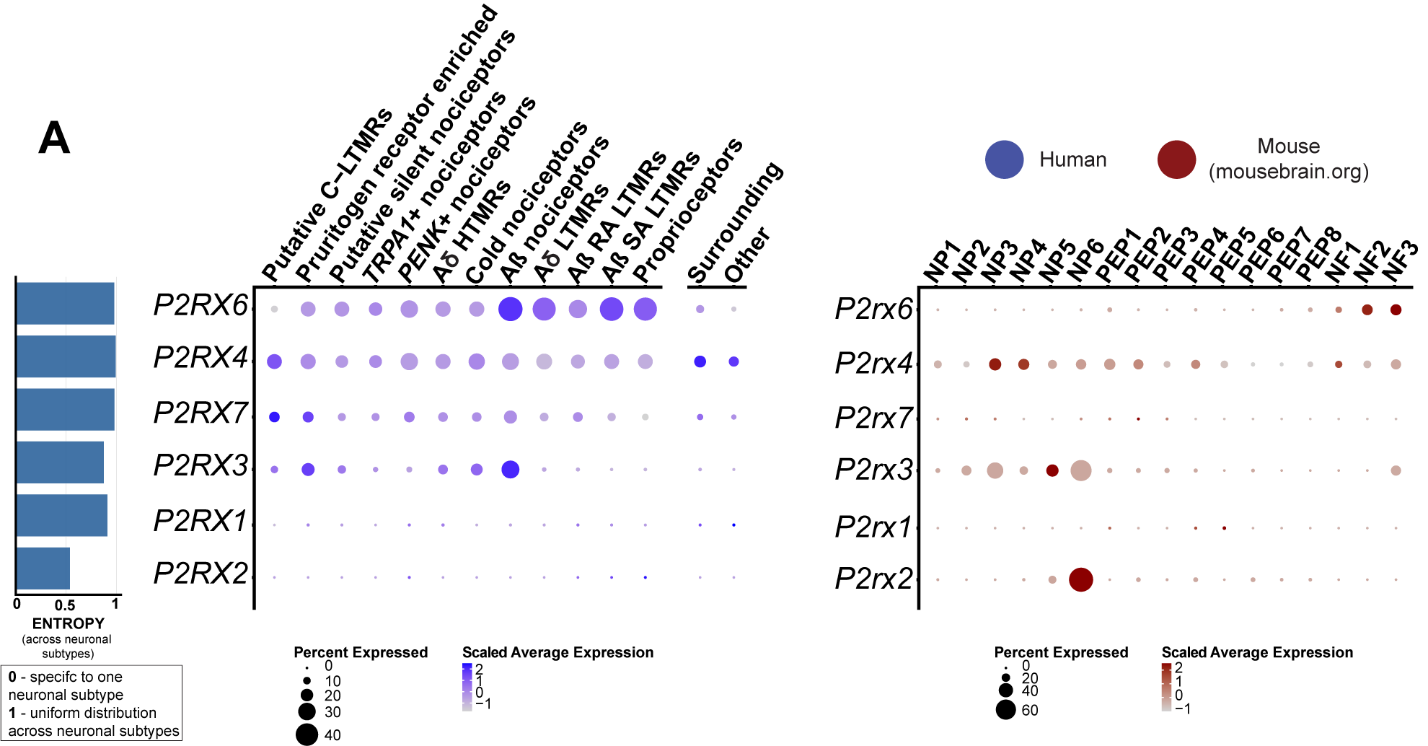

**Figure S22: Expression of ionotropic purinergic receptor genes in human and mouse datasets. (A)** Dot-plots showing the expression of purinergic receptor genes in human spatial transcriptomic (in blue) and mouse single-cell experiments (in red). The size of the dot represents the percentage of barcodes within a cluster and the color corresponds to the average expression (scaled data) across all barcodes within a cluster for each gene shown. Normalized entropy was used as a measure of “specificity of neuronal subtype”, where a score of 0 means a gene is specific to one neuronal subtype and 1 means that a gene has uniform distribution across neuronal subtypes.

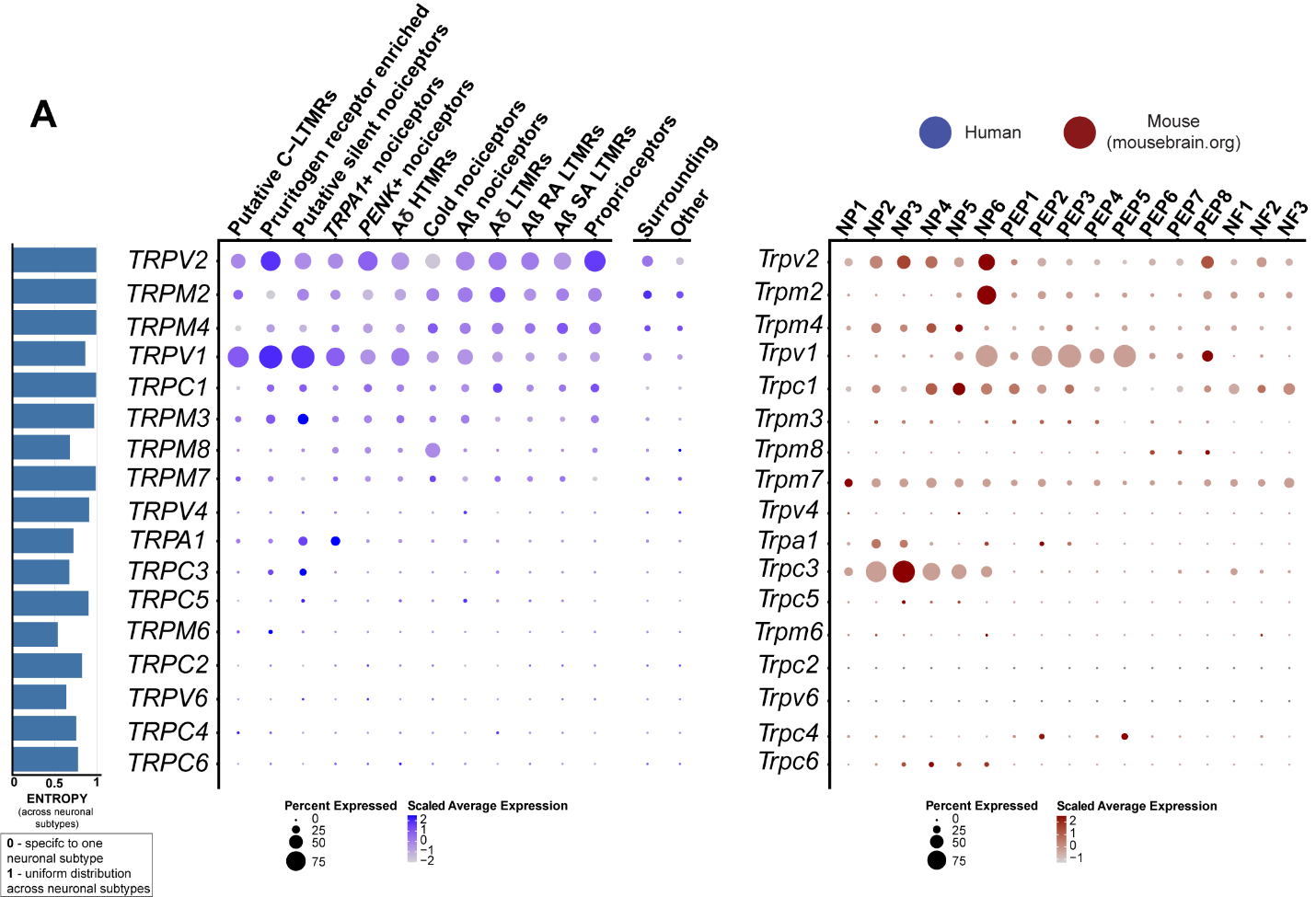

**Figure S23: Expression of transient receptor potential channel genes in human and mouse datasets. (A)** Dot-plots showing the expression of transient receptor potential genes in human spatial transcriptomic (in blue) and mouse single-cell experiments (in red). The size of the dot represents the percentage of barcodes within a cluster and the color corresponds to the average expression (scaled data) across all barcodes within a cluster for each gene shown. Normalized entropy was used as a measure of “specificity of neuronal subtype”, where a score of 0 means a gene is specific to one neuronal subtype and 1 means that a gene has uniform distribution across neuronal subtypes.

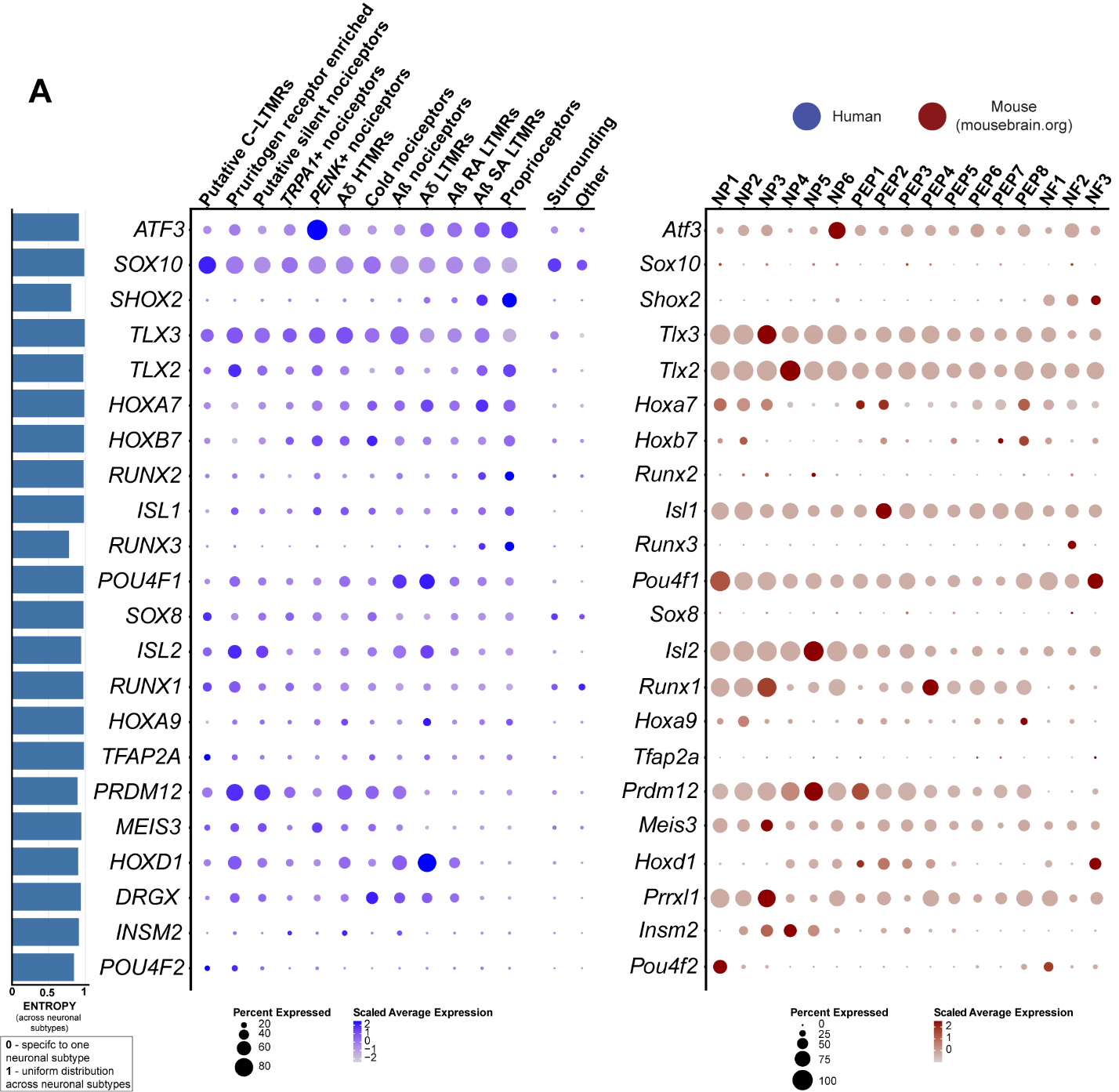

**Figure S24: Expression of neuronal transcription factors in human and mouse datasets. (A)** Dot-plots showing the gene expression of transcription factors involved in neuronal differentiation in human spatial transcriptomic (in blue) and mouse single-cell experiments (in red). The size of the dot represents the percentage of barcodes within a cluster and the color corresponds to the average expression (scaled data) across all barcodes within a cluster for each gene shown. Normalized entropy was used as a measure of “specificity of neuronal subtype”, where a score of 0 means a gene is specific to one neuronal subtype and 1 means that a gene has uniform distribution across neuronal subtypes.

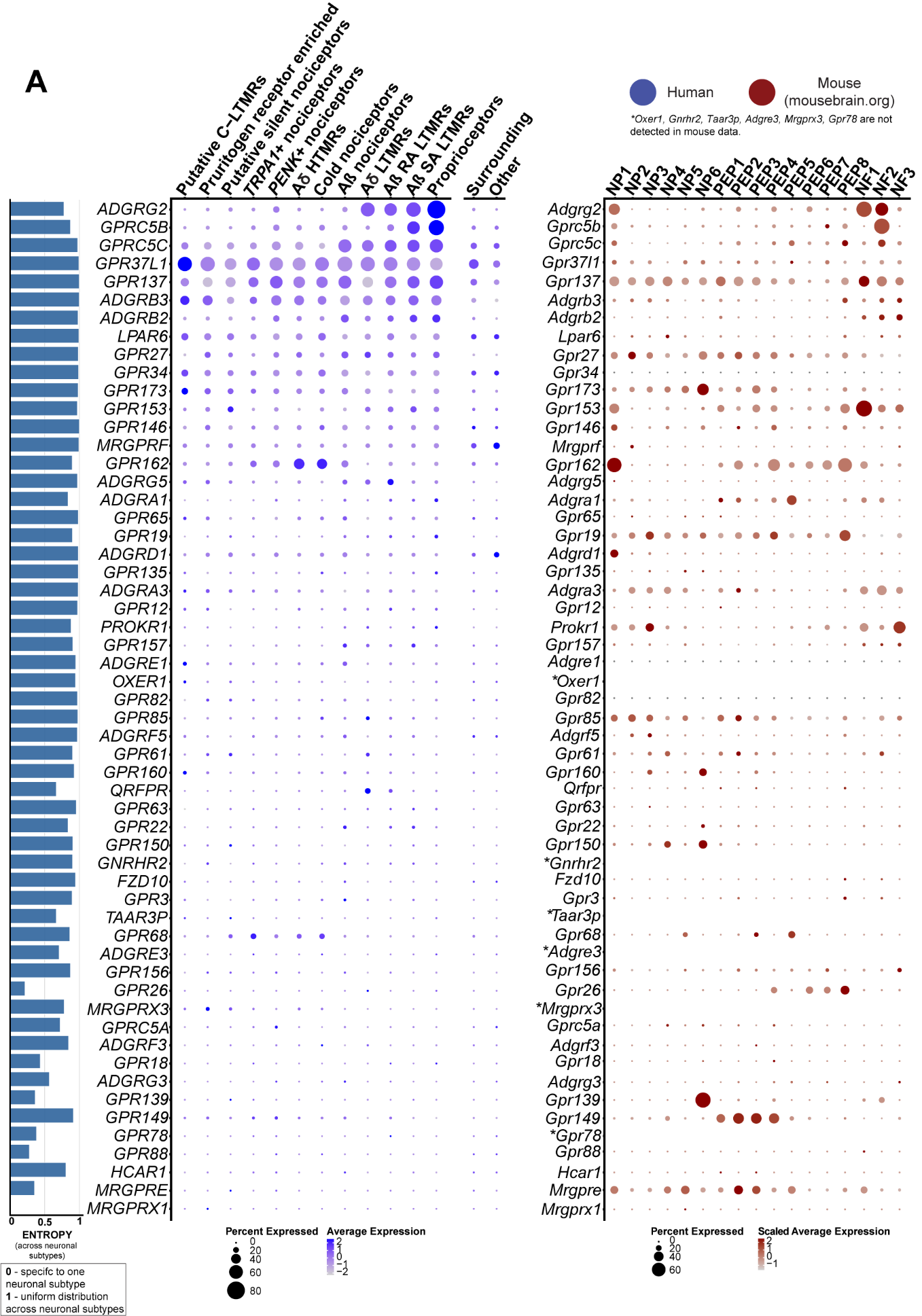

**Figure S25: Expression of understudied GPCRs in human and mouse datasets. (A)** Dot-plots showing the expression of understudied GPCR of the druggable genome in human spatial transcriptomic (in blue) and mouse single-cell experiments (in red). The size of the dot represents the percentage of barcodes within a cluster and the color corresponds to the average expression (scaled data) across all barcodes within a cluster for each gene shown. Normalized entropy was used as a measure of “specificity of neuronal subtype”, where a score of 0 means a gene is specific to one neuronal subtype and 1 means that a gene has uniform distribution across neuronal subtypes.

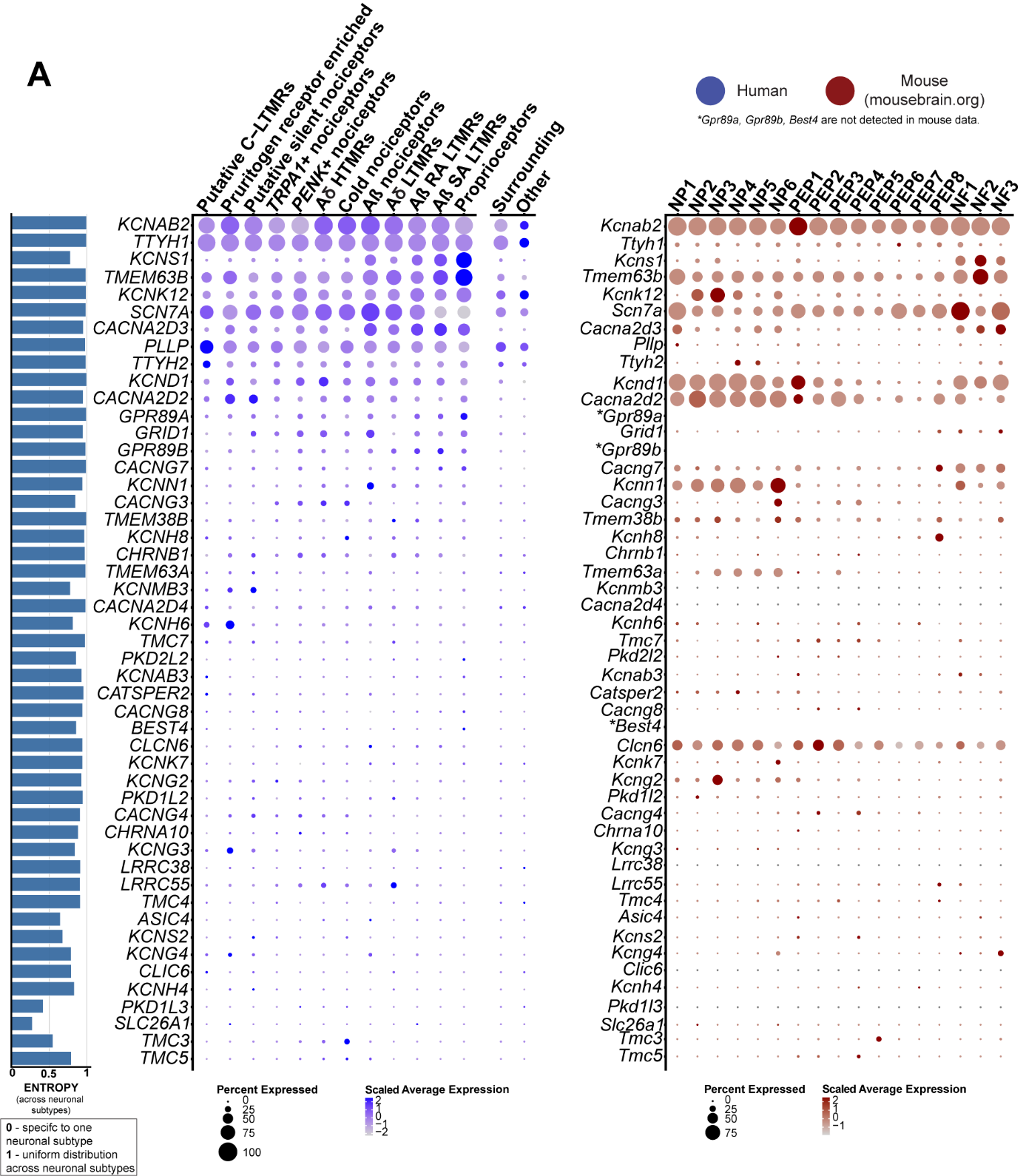

**Figure S26: Expression of** **understudied ion channels in human and mouse datasets. (A)** Dot-plots showing the expression of understudied ion channels of the druggable genome in human spatial transcriptomic (in blue) and mouse single-cell experiments (in red). The size of the dot represents the percentage of barcodes within a cluster and the color corresponds to the average expression (scaled data) across all barcodes within a cluster for each gene shown. Normalized entropy was used as a measure of “specificity of neuronal subtype”, where a score of 0 means a gene is specific to one neuronal subtype and 1 means that a gene has uniform distribution across neuronal subtypes.

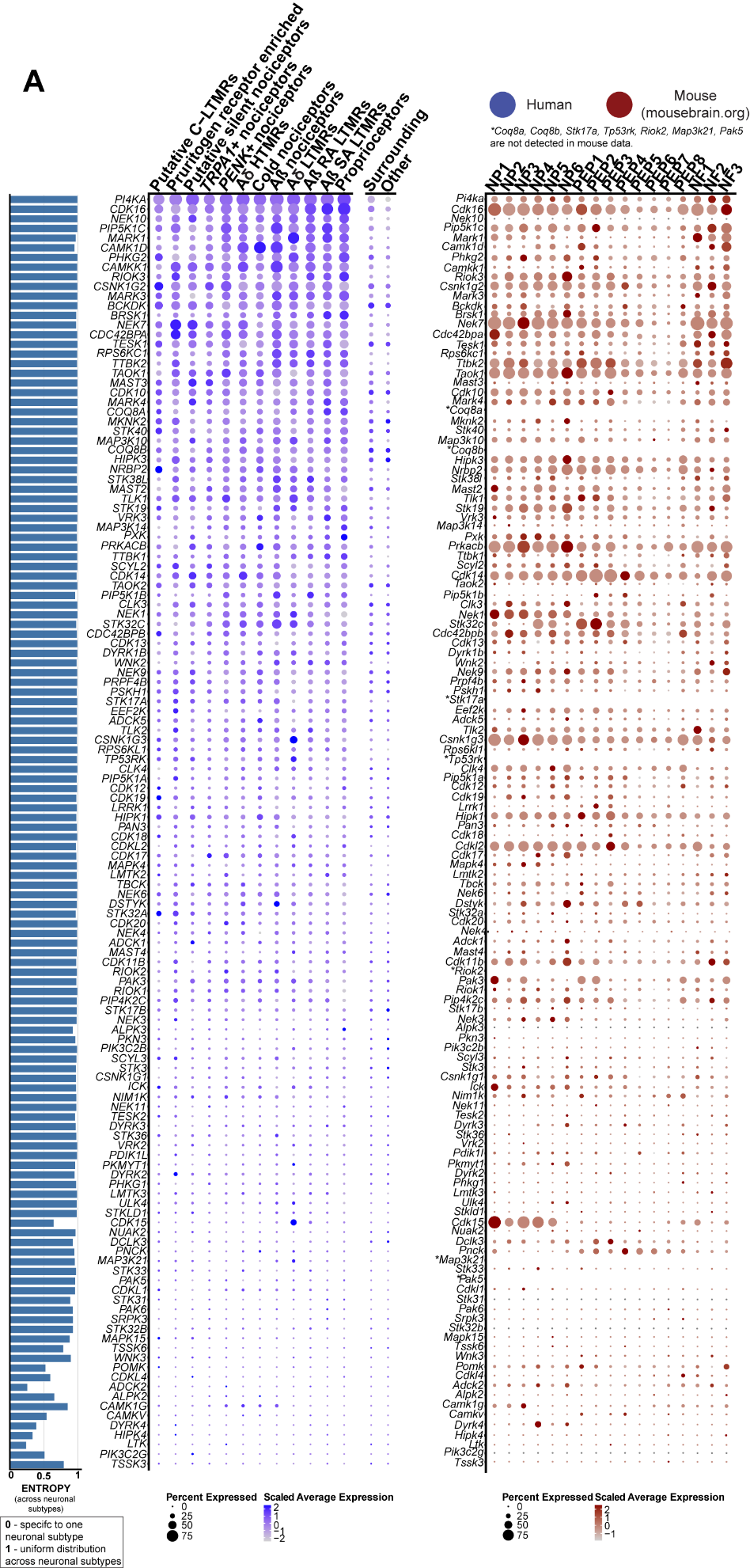

**Figure S27: Expression of understudied kinases in human and mouse datasets. (A)** Dot-plots showing the expression of understudied kinases of the druggable genome in human spatial transcriptomic (in blue) and mouse single-cell experiments (in red). The size of the dot represents the percentage of barcodes within a cluster and the color corresponds to the average expression (scaled data) across all barcodes within a cluster for each gene shown. Normalized entropy was used as a measure of “specificity of neuronal subtype”, where a score of 0 means a gene is specific to one neuronal subtype and 1 means that a gene has uniform distribution across neuronal subtypes.

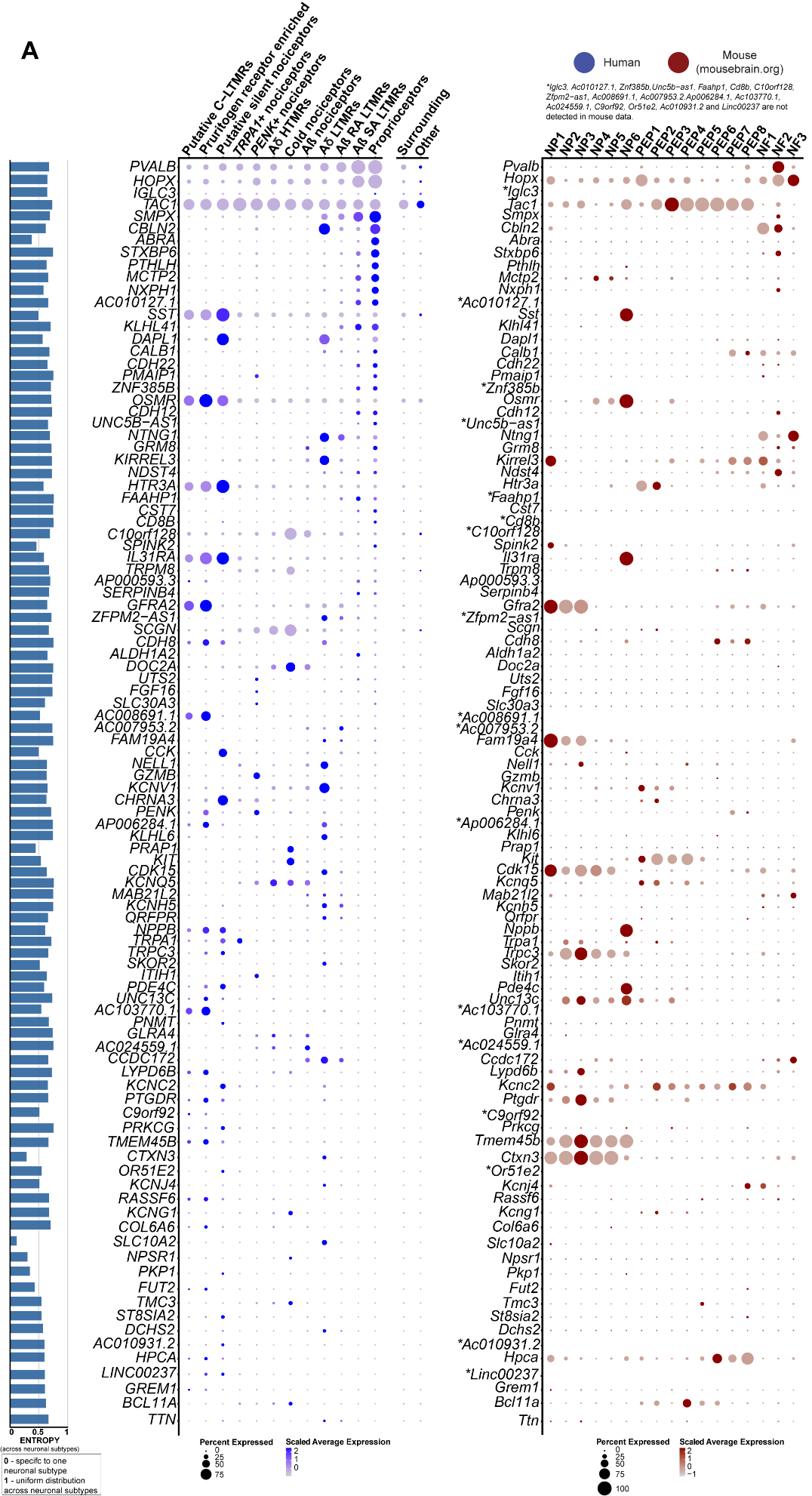

**Figure S28: Expression of low entropy genes. (A)** Dot-plots showing the expression of genes with low entropy from human Visium data. Human spatial transcriptomic data in blue and mouse single-cell experiments in red. The size of the dot represents the percentage of barcodes within a cluster and the color corresponds to the average expression (scaled data) across all barcodes within a cluster for each gene shown. Normalized entropy was used as a measure of “specificity of neuronal subtype”, where a score of 0 means a gene is specific to one neuronal subtype and 1 means that a gene has uniform distribution across neuronal subtypes.

**Table S1. Human donor information.**

| Donor # | Visium sample ID | Age | Sex | Race | Cause of death |
| --- | --- | --- | --- | --- | --- |
| 1 | F1 | 44 | F | Black | Anoxia/Cardiac Arrest |
| 2 | M1 | 29 | M | Black | Head Trauma |
| 3 | F2 | 57 | F | Black | Stroke |
| 4 | M2 | 53 | M | White | Head Trauma |
| 5 | F3 | 34 | F | White | Anoxia/Overdose |
| 6 | M3 | 65 | M | Black | Stroke |
| 7 | F4 | 53 | F | White | Stroke |
| 8 | M4 | 24 | M | White | Head Trauma |
| 9 | N/A | 56 | M | White | Stroke |
| 10 | N/A | 22 | M | Hispanic | Anoxia/Asphyxiation |
| 11 | N/A | 55 | M | White | Anoxia/Seizure |
| 12 | N/A | 48 | F | N/A | Head trauma |
| 13 | N/A | 18 | F | N/A | N/A |
| 14 | N/A | 61 | F | White | Anoxia/Cardiac Arrest |

**Table S2. Summary of RNAscope experiment details.**

| **mRNA** | **Experiment Combination** | **ACD Probe Cat No.** | **DRG Donor #** | **Total neurons assessed** |
| --- | --- | --- | --- | --- |
| *NPPB* | + *CALCA* | 448511 | 11-13 | 533 |
| *PRMD12* | + *CALCA* | 559961 | 9-10 | 368 |
| *SST* | + *CALCA* | 310591 | 5, 11-12 | 450 |
| ***CALCA*** |  | 605551-C2 | 5, 9-13 | **1351** |
| *NTRK1* | + *TRPV1* and *SCN10A* | 402631 | 2, 4-6 | 804 |
| *NTRK2* | + *TRPV1* and *SCN10A* | 402621 | 3, 5, 8 | 518 |
| *NTRK3* | + *TRPV1* and *SCN10A* | 406341 | 2, 7-8 | 649 |
| ***SCN10A*** |  | 406291-C2 | 2-8 | **1971** |
| ***TRPV1*** |  | 415381-C3 | 2-8 | **1971** |
| *PVALB* | +*LPAR3* and *TRPV1* | 422181-C3 | 1-3, 7 | 1161 |
| *LPAR3* | +*PVALB* and *TRPV1* | 428811-C1 | 1-3, 7 | 1161 |
| *TRPV1* | +*PVALB* and *LPAR3* | 415381-C2 | 1-3, 7 | 1161 |
| *PENK* | +*TRPM8* | 548301-C3 | 5-6, 8, 14 | 965 |
| *TRPM8* | +*PENK* | 543121-C1 | 5-6, 8, 14 | 965 |

**File S1. (separate Excel file)**

Ranked gene expression within each neuronal barcode by cluster.

**Files S2-27. (separate Excel files)**

Results of statistical analysis for sex differences. (A) Results for overall genes tested. (B) Genes differentially expressed. (C) Genes differentially expressed specifically in neurons (only Files S2-14). Files S2-14 are files for neuronal barcodes and files S15-27 are for surrounding barcodes.

**File S28. (separate Excel file)**

Gene enrichment analysis for the genes DE in pruritogen receptor enriched population.

**File S29. (separate Excel file)**

Overall sequencing statistics for Visium experiments.

**File S30. (separate Excel file)**

Data underlying human and macaque DRG gene expression comparison.
